# Mural cell contractility regulates vessel diameter by controlling cell morphology and vessel coverage

**DOI:** 10.64898/2026.06.07.730635

**Authors:** Mingzhao Hu, Hedele Zeng, Ricard Casanovas-Zapata, Koji Ando, Yukiko T. Matsunaga, Li-Kun Phng

## Abstract

Mural cells regulate vascular architecture, but how their contractility and morphology together determine vessel diameter is unclear. Using high-resolution live imaging and single-cell morphometrics in zebrafish, we show that vSMCs and pericytes progressively retract actin-rich processes and decrease in size during development. Mural cell-specific manipulation of RhoA activity altered cell morphology and vessel coverage, revealing vessel- and stage-dependent effects on vascular calibre. vSMC contractility is dispensable for initial dorsal aorta narrowing but is required to maintain its constricted diameter and to limit vessel wall deformation during pulsatile haemodynamic stimulation. By contrast, pericyte contractility is dispensable for constriction and stabilisation of intersegmental vessels. In cerebral vessels, both increased and reduced RhoA activity caused larval dilation, whereas elevated RhoA activity is associated with focal narrowing and stenosis in adults. Thus, mural cell contractility regulates vascular calibre and mechanics not through force alone, but through its integrated effects on cell morphology and vessel coverage.

## Introduction

Blood vessels are complex multicellular tubes lined by endothelial cells and supported along their abluminal surface by mural cells, which include vascular smooth muscle cells (vSMCs) and pericytes^1^. vSMCs primarily surround arteries and arterioles, whereas pericytes associate predominantly with capillaries and smaller vessels^2^.

Mural cells originate from heterogeneous progenitor populations, including mesenchymal and neural crest–derived cells, and are recruited to nascent vessels primarily through PDGF-B/PDGFRβ signalling^1–4^. During development, mural cells undergo progressive differentiation into distinct subtypes along the vascular tree. vSMCs acquire a contractile phenotype characterised by high expression of smooth muscle markers such as α-smooth muscle actin (Acta2) and transgelin (Tagln), enabling efficient force generation around larger vessels^5,6^. In contrast, pericytes are characterised by lower and more variable expression of classical smooth muscle contractile markers—including Acta2, smooth muscle myosin heavy chain (MYH11), calponin (CNN1) and smoothelin (SMTN)—compared to vSMCs, reflecting their less mature contractile state^7,8^. Increasing evidence suggests that mural cells span a continuous spectrum of morphological, molecular and functional states, forming a hierarchical and structurally diverse network rather than discrete cell populations, with intermediate phenotypes observed in transitional vessel segments^8^.

Mural cells play diverse roles in vascular development and maintenance. During angiogenesis and vascular remodelling, mural cells coordinate endothelial sprouting, vessel maturation and pruning, thereby shaping vascular architecture^1,2,9^. During vascular remodelling, blood vessels undergo coordinated changes in diameter, structure and cellular composition to establish a functional and hierarchically organised network^10^. In zebrafish (*Danio rerio*), this process is well characterised, with major vessels such as the dorsal aorta (DA) undergoing an initial phase of dilation followed by progressive narrowing between 1-3 days post fertilisation (dpf), and intersegmental vessels (ISVs) undergoing gradual reduction in diameter between 2-4 dpf^11,12^. These changes have been widely attributed to endothelial cell–intrinsic mechanisms^10^. However, mural cells—including vSMCs and pericytes—are progressively recruited to nascent vessels during this period and are thought to contribute to vessel diameter regulation and stabilisation. vSMCs are specialised for contractile function and are the primary regulators of arterial tone, controlling vessel diameter and blood flow^13^. In addition to this canonical role, vSMCs contribute to vessel stabilisation by providing mechanical support and promoting extracellular matrix deposition^14,15^. In contrast, pericytes exhibit more heterogeneous and context-dependent functions^2,16,17^. They are essential for maintaining endothelial barrier integrity, particularly within the blood–brain barrier, and regulate endothelial cell behaviour through paracrine signalling pathways^7^. While considered less contractile than vSMCs, accumulating evidence suggests that pericytes can modulate capillary diameter and local blood flow in certain tissues and developmental stages^16–18^, although the extent and physiological significance of this contractility remain debated^13^.

Mural cells exhibit diverse shapes. vSMCs typically exhibit a compact morphology with circumferential processes that wrap around vessels, whereas pericytes display elongated cell bodies with longitudinal and branching processes that extend along and partially encircle the vessel wall^19–21^. Notably, pericytes comprise multiple morphological subtypes that vary across vascular beds and developmental or physiological contexts. In the adult mouse cerebral cortex, ensheathing pericytes on pre-capillary arterioles exhibit short primary processes and broad secondary processes that nearly encircle the vessel^8,21,22^, whereas thin-strand pericytes on capillaries possess long, thin longitudinal processes with sparse circumferential coverage^8,13^. Stellate pericytes, associated with post-capillary venules, display a more complex, radially branching morphology with irregular, mesh-like processes^13,23^. Stellate-shaped pericytes with extensive protrusions have also been described on angiogenic blood vessels of the early postnatal mouse retina^23^. These distinct morphologies likely underlie specialised modes of interaction with endothelial cells and differential regulation of vascular function^22^. Despite this morphological diversity, transcriptomic analyses of brain vasculature indicate that pericytes across distinct vascular segments exhibit highly similar gene expression profiles and are often transcriptionally indistinguishable^24,25^. This suggests that mural cell morphology is not solely specified by transcriptional identity but is also shaped by context-dependent signalling mechanisms, including regulation by the actin cytoskeleton and small Rho GTPases^26^. RhoA activity is associated with actomyosin contractility and cell body constriction, whereas the related GTPase Cdc42 promotes the formation of filopodia-like protrusions and cellular extensions. Consistent with this, Cdc42 has been shown to be essential for mural cell migration, proliferation and patterning in the postnatal mouse retina, where it drives the formation of protrusive structures in pericytes during sprouting angiogenesis^27^. In contrast, elevated RhoA/ROCK signalling and contractility in α-parvin-deficient vSMCs induce a more rounded morphology and impaired cell spreading^28^. Together, these observations suggest that distinct cytoskeletal and signalling programmes, rather than transcriptional differences alone, govern mural cell shape and behaviour.

At present, little is known about mural cell shape dynamics during vascular remodelling, and how these contribute to vessel diameter control. It remains unclear how mural cell contractility regulates vessel diameter, and whether this occurs through direct vessel constriction or indirectly by reshaping cell morphology and vessel coverage. The zebrafish model offers unique advantages for understanding these relationships *in vivo*. Its optical transparency enables high-resolution live imaging of vascular remodelling, allowing direct measurement of vessel diameter changes alongside dynamic observation of mural cell shape^29^. In this study, we combine transgenic labelling, single-cell morphometric analysis and genetic manipulation to define the relationship between mural cell contractility, morphology and vessel diameter during vascular remodelling.

## Results

### Enhanced visualisation of mural cell morphology using Lifeact

The *TgBAC(pdgfrb:EGFP)^ncv^*^22^ transgenic zebrafish line (hereafter referred to as *PDGFRβ–EGFP*) is widely used to visualise mural cells in zebrafish and broadly labels mural cell bodies and their elongated processes^20^. However, because EGFP is distributed throughout the cytoplasm, thin cellular processes are often poorly resolved. To improve the visualisation of actin-rich structures, we employed a GAL4/UAS-based strategy to drive mural cell–specific expression of Lifeact–mCherry under the control of the *pdgfrb* promoter (Supp. Fig. 1A). Combined with high-resolution live imaging, Lifeact–mCherry expression enabled clear visualisation of thin cellular processes that are not readily resolved in *PDGFRβ–EGFP* cells (Supp. Fig. 1B, C). Using this labelling strategy, we established a systematic and unbiased analytical framework to quantify mural cell morphometrics^30^ (Supp. Fig. 2) from 3 to 14 dpf, a period characterised by active vascular remodelling.

**Figure 1.**
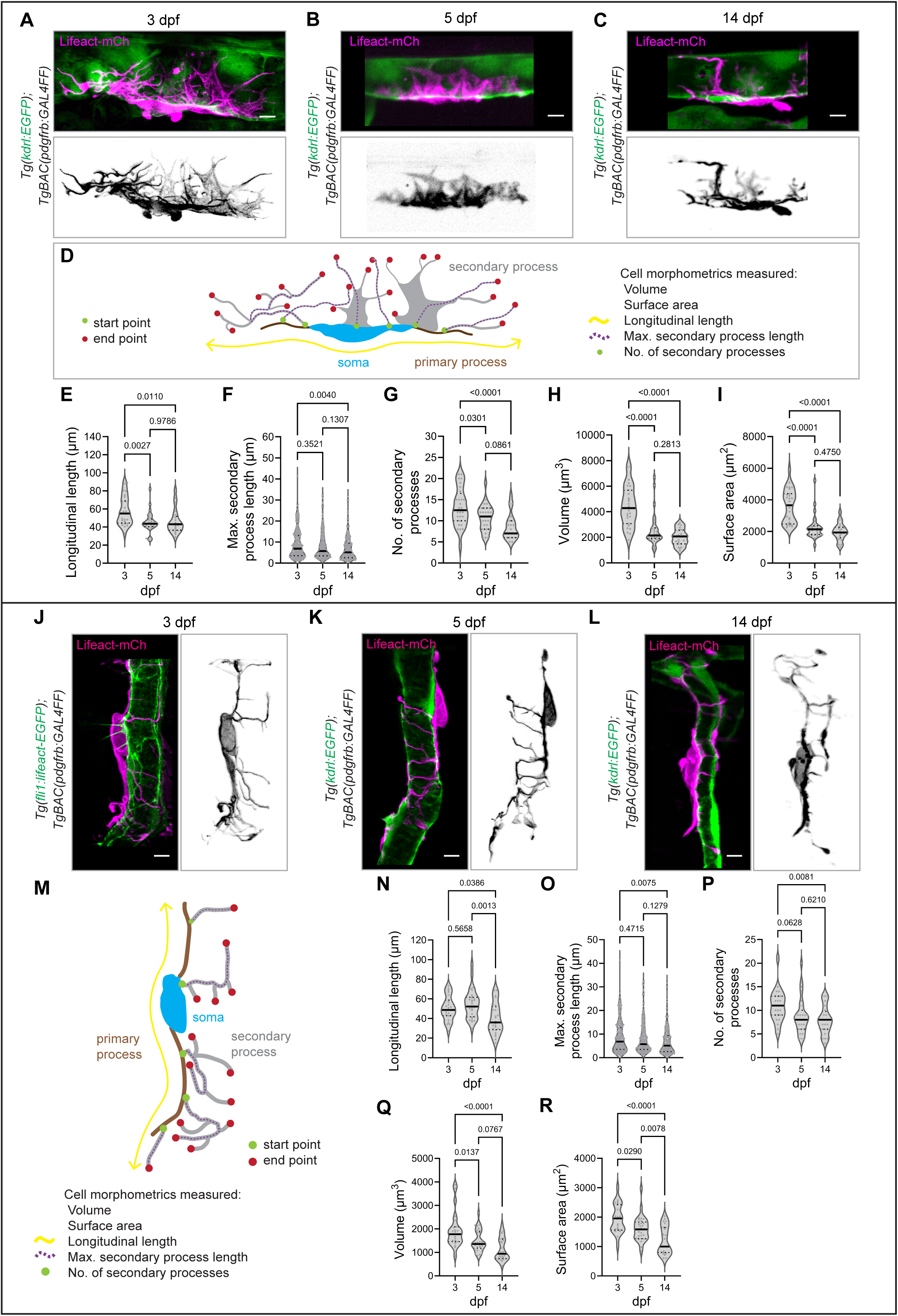
Trunk mural cells progressively reduce cellular processes and size during development. **A – C** Maximum intensity projection of confocal z-stacks showing vSMCs expressing Lifeact-mCherry along the DA at 3, 5 and 14 dpf in *TgBAC(pdgfrb:GAL4FF); Tg(kdrl:EGFP)* embryos. **D** Schematic illustrating morphometric parameters measured for vSMC analysis. Start points (green) denote the origin of each secondary process, and the total number of start points defines the number of secondary processes. End points (red) mark the distal terminal of secondary processes. The maximum secondary process length is defined as the longest distance between a given start point and all corresponding end points arising from that start point. The longitudinal length (yellow line) is defined as the vessel-length distance between the two distal ends of the vSMC. Cell volume and surface area were measured across the stages. **E – I** Quantification of vSMC longitudinal length, maximum secondary process length, number of secondary processes, cell volume and surface area at 3, 5 and 14 dpf (n = 27 vSMCs from 12 embryos at 3 dpf; n = 29 vSMCs from 13 embryos at 5 dpf; n = 21 vSMCs from 6 embryos at 14 dpf). Data are collected from 2 (3 dpf), 2 (5 dpf) and 2 (14 dpf) independent experiments and analyzed by ordinary one-way ANOVA with Tukey’s multiple comparisons test. **J – L** Maximum intensity projection of confocal z-stacks showing pericytes expressing Lifeact-mCherry associated with ISVs at 3, 5 and 14 dpf in *TgBAC(pdgfrb:GAL4FF); Tg(fli1:Lifeact-EGFP)* or *TgBAC(pdgfrb:GAL4FF); Tg(kdrl:EGFP)* embryos. **M** Schematic illustrating morphometric parameters used for pericyte analysis. **N – R** Quantification of pericyte longitudinal length, maximum secondary process length, number of secondary processes, cell volume and surface area at 3, 5 and 14 dpf (n = 23 pericytes from 12 embryos at 3 dpf; n = 29 pericytes from 17 embryos at 5 dpf; n = 24 pericytes from 10 embryos at 14 dpf). Data were collected from 3 (3 dpf), 2 (5 dpf) and 2 (14 dpf) independent experiments and analysed by ordinary one-way ANOVA with Tukey’s multiple comparisons test. Violin plots represent the entire range of values, dotted lines indicate first and third quartiles, center lines are median. Scale bar, 5 µm.

**Figure 2.**
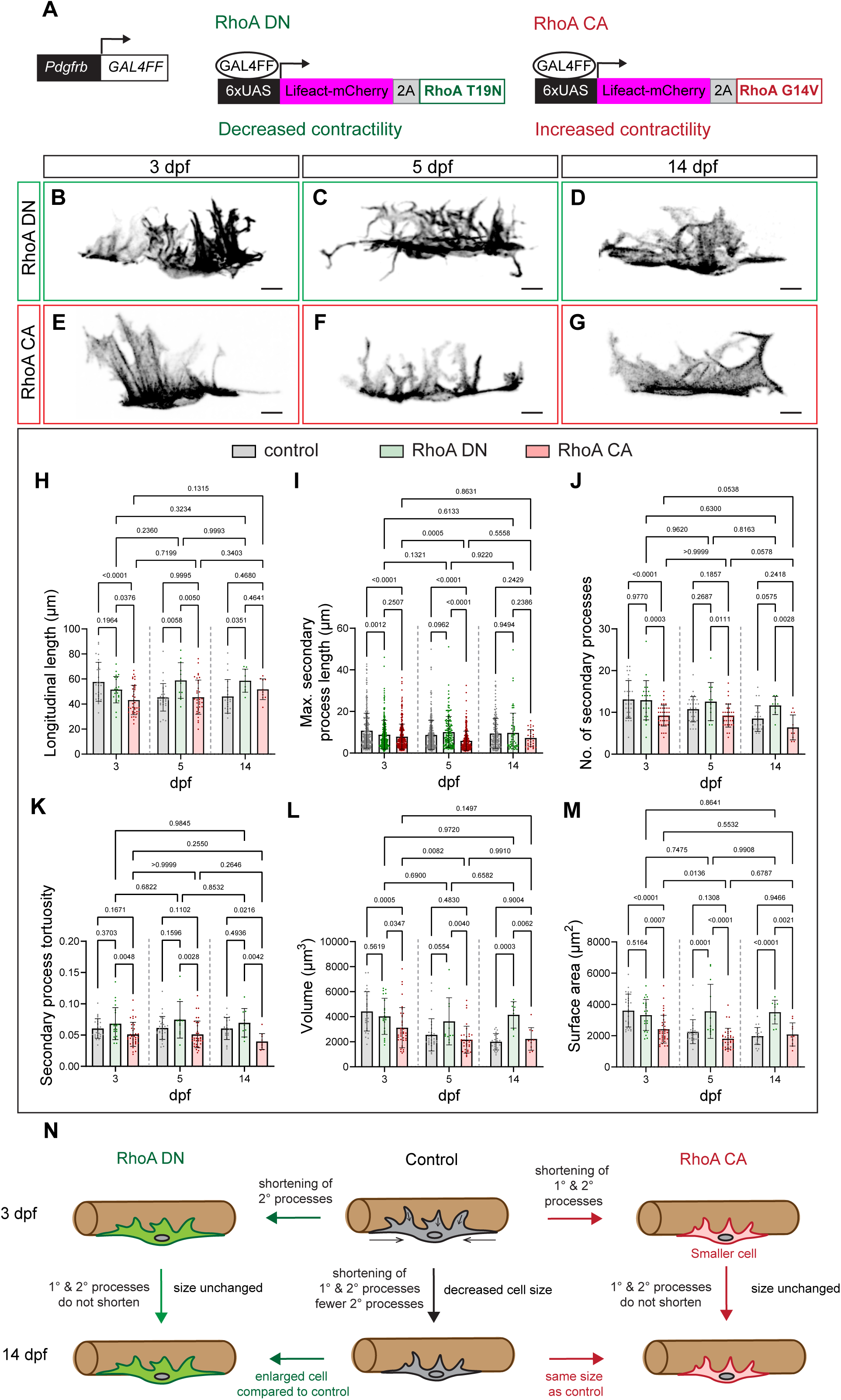
Actomyosin contractility drives vSMC process retraction and compaction. **A** Plasmid constructs encoding GAL4FF/UAS-driven expression of dominant-negative RhoA T19N or constitutively active RhoA G14V linked via a self-cleaving P2A peptide fused to Lifeact-mCherry. **B – G** Maximum intensity projection of confocal z-stacks of vSMCs expressing RhoA DN (**B – D**) or RhoA CA (**E – G**) at 3, 5 and 14 dpf embryos. Scale bar, 5 µm. **H – M** Quantification of vSMC longitudinal length, maximum secondary process length, number of secondary processes, tortuosity of secondary processes, cell volume and surface area at 3, 5 and 14 dpf (RhoA DN: n = 22 vSMCs from 13 embryos at 3 dpf; n = 11 vSMCs from 7 embryos at 5 dpf; n = 9 vSMCs from 6 embryos at 14 dpf; RhoA CA: n = 38 vSMCs from 17 embryos at 3 dpf; n = 34 vSMCs from 19 embryos at 5 dpf; n = 10 vSMCs from 8 embryos at 14 dpf from 3 (3 dpf), 2 (5 dpf) and 2 (14 dpf) independent experiments). Data were analysed by two-way ANOVA with Tukey’s multiple comparisons test. **N** Schematic illustration of trunk vSMC morphological changes during vascular remodelling and the effects of perturbed RhoA signalling on vSMC shape.

### Trunk mural cells decrease in cellular process length and size during development

To distinguish mural cells from other *pdgfrb*-expressing perivascular cells in the zebrafish trunk, we identified mural cells by their high *PDGFRβ–EGFP* reporter intensity and morphology, namely elongated processes that closely extend along or wrap around the vessel wall (Supp. Fig. 1B, C). We further distinguished vSMCs from pericytes according to their vascular location and previously characterised molecular identities. Mural cells surrounding the DA express the vSMCs genes, *acta2* and *tagln*^5,31,32^ but lack the expression of the pericyte genes, *ndufa4l2a* and *abcc9*^33^. In contrast, mural cells associated with arterial and venous ISVs (aISVs and vISVs) express *ndufa4l2a* and *abcc9*^33^ but lack *acta2* and *tagln* expression^5^. We therefore refer to *PDGFRβ–EGFP*^high^ expressing cells wrapping the DA as vSMCs while those with processes closely associated with aISVs and vISVs as pericytes.

vSMCs lining the ventral floor of the DA extend primary processes along the longitudinal axis of the vessel and secondary processes that project circumferentially across the vessel wall (Fig. 1A–D). These secondary processes comprise actin-rich bundles interspersed with lamellar protrusions that extend across the DA in a tortuous manner. Between 3 and 5 dpf, longitudinal length decreased significantly and subsequently remained stable until 14 dpf (Fig. 1E). Similarly, the maximum length of secondary processes shows a decreasing trend over time, with a significant reduction between 3 and 14 dpf (Fig. 1F). In parallel, the number of secondary processes (Fig. 1G) reduces over this period. As a result, vSMCs significantly reduce in cell volume (Fig. 1H) and surface area (Fig. 1I) between 3 and 5 dpf and remain largely unchanged thereafter.

Pericytes associated with ISVs typically exhibit a protruding cell body that extend long, slender primary processes bilaterally along the longitudinal axis of the vessel and circumferential secondary processes that do not encircle the vessel (Fig. 1J–M). Morphometric analysis revealed that pericyte longitudinal length (Fig. 1N), maximum secondary process length (Fig. 1O) and the number of secondary processes (Fig. 1P) are significantly reduced at 14 dpf compared with 3 dpf. This shortening of secondary processes is accompanied by significant reductions in cell volume (Fig. 1Q) and surface area (Fig. 1R) by 14 dpf.

In sum, trunk mural cells shrink during development through progressive process shortening. vSMCs undergo pronounced compaction between 3 and 5 dpf and remain compact through 14 dpf, whereas pericytes show a more gradual reduction in process length and cell size between 3 and 14 dpf.

### RhoA–actomyosin contractility drives vSMC process retraction and compaction

The reduction in vSMC and pericyte size during zebrafish development suggests that intrinsic contractility drives cytoskeletal reorganisation and membrane retraction to reshape mural cells. Analysis of publicly available single-cell transcriptomic data^31^ revealed that vSMCs and pericytes express genes encoding non-muscle myosin II (*myh9b* and *myh10*), smooth muscle myosin (*myh11a*), myosin regulatory light chain (*myl9a*, *myl9b*, *myl12.1* and *myl12.2*) and the GTPase RhoA (*rhoaa*, *rhoab* and *rhoac,* Supp. Fig. 3A). Imaging of Myl9b-mCherry, previously used to observe the localisation and dynamics of non-muscle myosin II associated along actin cables in zebrafish endothelial cells^11,34^, revealed its localisation in the actin-rich primary and secondary processes of vSMCs and pericytes (Supp. Fig. 3B, C). Together, these expression and localisation data indicate that mural cells possess a contractile machinery capable of driving morphological changes.

**Figure 3.**
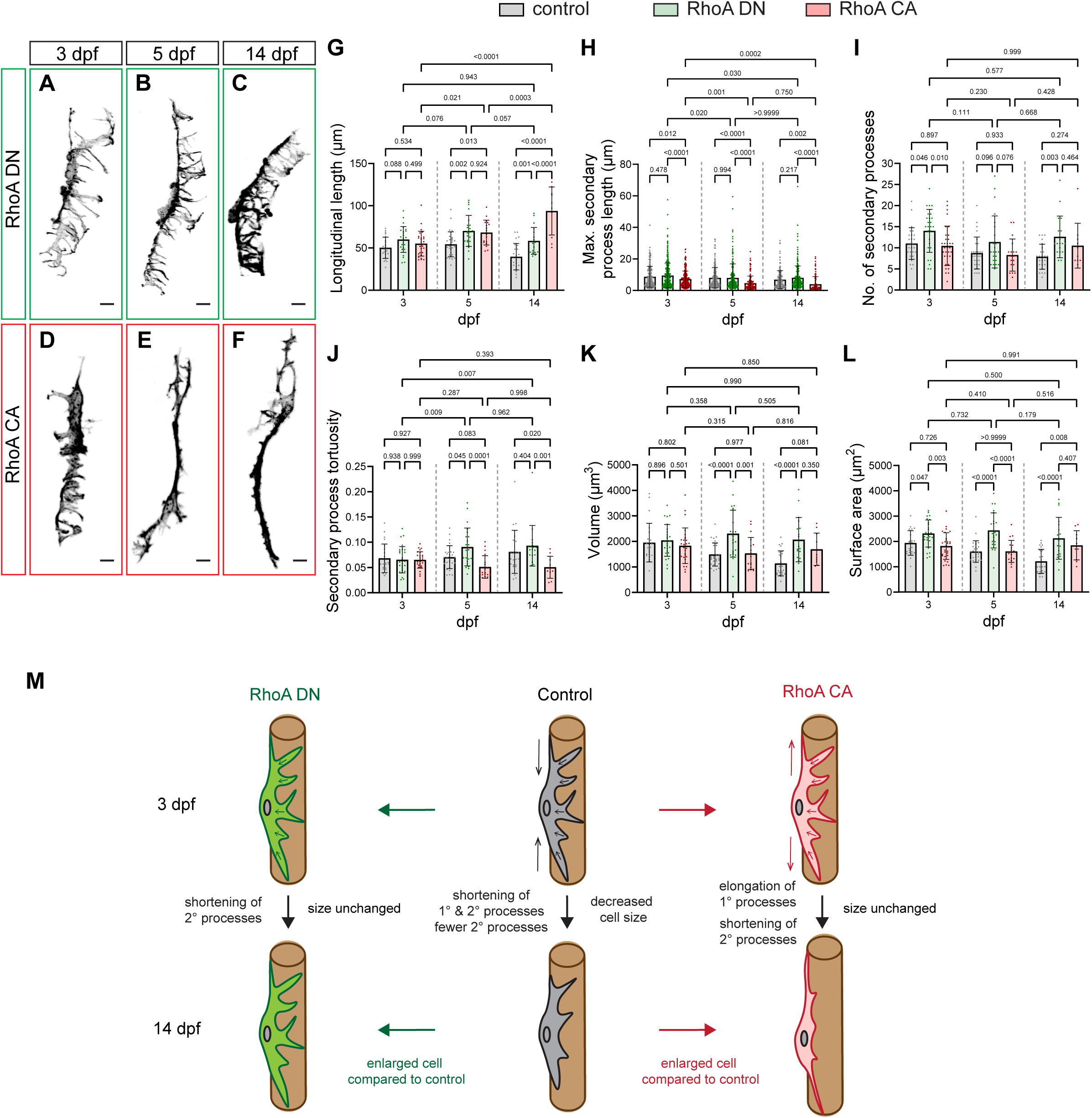
Balanced actomyosin activity shapes pericyte morphology. **A – F** Maximum intensity projection of confocal z-stacks of pericytes expressing RhoA DN (**A – C**) or RhoA CA (**D – F**) at 3, 5 and 14 dpf embryos. Scale bar, 5 µm. **G - L** Quantification of pericyte longitudinal length, maximum secondary process length, number of secondary processes, tortuosity of secondary processes, cell volume and surface area at 3, 5 and 14 dpf (RhoA DN: n = 26 pericytes from 15 embryos at 3 dpf; n = 23 pericytes from 16 embryos at 5 dpf; n = 18 pericytes from 7 embryos at 14 dpf; RhoA CA: n = 28 pericytes from 15 embryos at 3 dpf; n = 18 pericytes from 11 embryos at 5 dpf; n = 10 pericytes from 6 embryos at 14 dpf from 3 (3 dpf), 2 (5 dpf) and 2 (14 dpf) independent experiments.) Data were analysed by two-way ANOVA with Tukey’s multiple comparisons test. **M** Schematic illustration of trunk pericyte morphological changes during vascular remodelling and the effects of perturbed RhoA signalling on pericyte shape.

Because RhoA–ROCK signalling promotes actin assembly and myosin light-chain phosphorylation to enhance mural cell contractility ^35–37^, we tested whether intrinsic contractility drives membrane retraction and shape transitions by manipulating RhoA activity in individual mural cells. Using the GAL4/UAS system, we expressed either dominant negative RhoA T19N (hereafter RhoA DN) or constitutively active RhoA G14V (hereafter RhoA CA), each linked via a self-cleaving P2A peptide to Lifeact–mCherry (6×UAS:RhoA-T19N-P2A-Lifeact–mCherry or 6×UAS:RhoA-G14V-P2A-Lifeact–mCherry), in *TgBAC(pdgfrb:GAL4FF)* zebrafish (Fig. 2A - G). This strategy enabled simultaneous manipulation of actomyosin contractility and visualisation of single-cell morphology.

In contrast to control vSMCs, vSMCs overexpressing RhoA DN failed to undergo shortening of longitudinal length and maximum secondary process length between 3 and 14 dpf (Fig. 2H, I). The number of secondary processes was comparable between RhoA DN-expressing and control vSMCs at 3 and 5 dpf but showed a trend towards an increase in RhoA DN-expressing cells at 14 dpf (Fig. 2J). Secondary process tortuosity was comparable between RhoA DN-expressing and control vSMCs at all developmental stages (Fig. 2K). As a result, RhoA DN–expressing vSMCs maintained cell volume (Fig. 2L) and surface area (Fig. 2M), rather than undergoing the developmental shrinkage observed in control cells (Fig. 1G, H).

Conversely, overexpression of RhoA CA in vSMCs resulted in pronounced morphological changes. At 3 dpf, these cells exhibited significantly reduced longitudinal length (Fig. 2H) and shorter secondary processes (Fig. 2I) compared with control cells. Secondary process length continued to decrease between 3 and 5 dpf before stabilising thereafter. In addition, the number of secondary processes was significantly reduced at 3, 5 and 14 dpf (Fig. 2J), indicating decreased branching complexity. Secondary processes in RhoA CA-expressing vSMCs were also significantly less tortuous than those in control cells at all developmental stages (Fig. 2K), consistent with their visibly straighter morphology (Fig. 2E–G). This early shortening of primary and secondary processes resulted in significantly reduced cell volume (Fig. 2L) and surface area (Fig. 2M) at 3 dpf, with no further size reduction thereafter, suggesting that vSMCs reached their minimal cell size earlier during development.

To independently reduce actomyosin contractility, we expressed phosphorylation-deficient Myl9b A2A3 together with Lifeact–mCherry in mural cells (6×UAS:Myl9b-A2A3-P2A-Lifeact–mCherry; Supp. Fig. 4A - C). At 5 dpf, Myl9b A2A3-expressing vSMCs showed increased longitudinal length, secondary processes, cell volume and surface area, while maximum secondary process length remained unchanged (Supp. Fig. 4D–H). Thus, reducing myosin II activity promotes a larger, more branched vSMC morphology, consistent with reduced RhoA activity.

**Figure 4.**
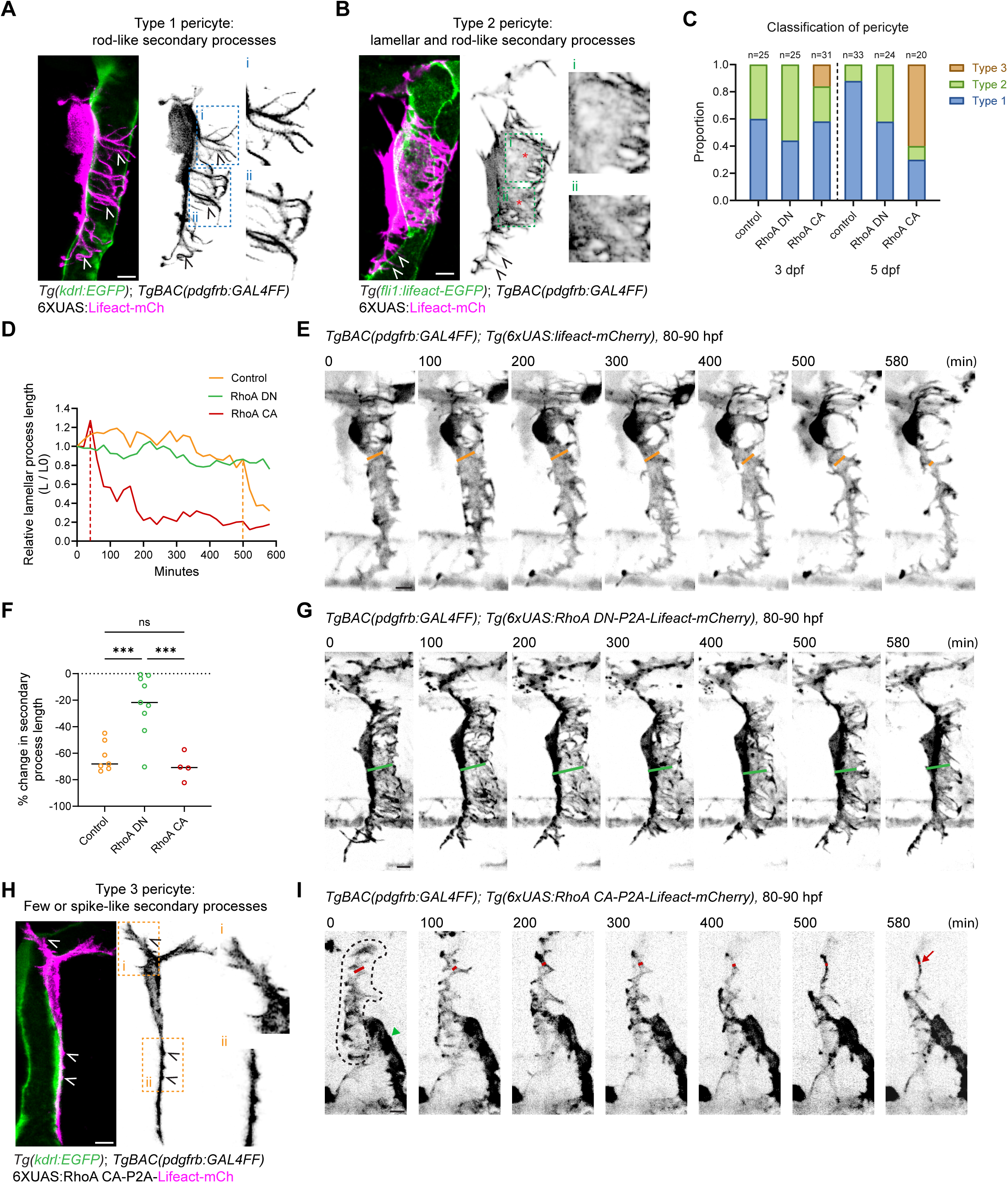
Retraction of pericyte secondary processes is actomyosin-dependent. **A - B** Maximum intensity projection of confocal z-stacks of Type 1 pericytes (**A**) and Type 2 pericytes (**B**) in control embryos at 3 dpf. Arrowheads indicate rod-like processes. Asterisks indicate lamellar structures. Insets show higher magnification views. **C** Proportion of each pericyte subtype at 3 and 5 dpf under control, RhoA DN and RhoA CA conditions (control: from 12/17 embryos at 3/5 dpf; RhoA DN: from 15/16 embryos at 3/5 dpf; RhoA CA: from 15/11 embryos at 3/5 dpf). Total number of pericytes for each condition is indicated on the top of the bar (at 3 dpf, control vs RhoA DN, Type 1 vs other types, Fisher’s exact test, p = 0.3961; at 5 dpf, control vs RhoA CA, Type 3 vs other types, Fisher’s exact test, p<0.0001). **D - I** RhoA activity modulates secondary process retraction. Change in relative lamellar secondary process length (**D**) and percentage change in secondary process length (**F**) between 80 and 90 hpf in control, RhoA DN- and RhoA CA-overexpressing pericytes (control: 7 pericytes from 4 embryos; RhoA DN: 9 pericytes from 5 embryos; RhoA CA: 4 pericytes from 3 embryos). Data were collected from 2 independent experiments and analysed by ordinary one-way ANOVA with Tukey’s multiple comparisons test. Center lines are median. Representative time-lapse images of control (**E**, from Supp. Video 1), RhoA DN-overexpressing (**G**, from Supp. Video 2) and RhoA CA-overexpressing (**I**, from Supp. Video 3) pericytes. Dashed outline in I indicates pericytes. Green arrowhead indicates perivascular fibroblast. Maximum intensity projection of confocal z-stacks of Type 3 pericytes induced by RhoA CA-overexpression (**H**). Arrowheads indicate spike-like processes. Insets show higher magnification views. Scale bar, 5 µm

Overall, these findings show that RhoA–actomyosin contractility drives developmental vSMC compaction. Reduced RhoA or myosin II activity maintains a larger, more branched morphology, whereas elevated RhoA activity accelerates process retraction and cell shrinkage.

### Balanced RhoA activity shapes pericyte morphology

Similar to vSMCs, RhoA DN-expressing pericytes failed to shorten longitudinally and remained longer than controls (Fig. 3A-C, G). They also formed significantly more secondary processes at 3 and 14 dpf, while secondary process length and tortuosity remained comparable to controls at each stage examined (Fig. 3H - J). Consequently, these cells failed to undergo developmental shrinkage and had larger volume and surface area at 5 and 14 dpf (Fig. 3K, L). Similarly, reducing myosin II activity through expression of Myl9b A2A3 increased pericyte size. At 5 dpf, Myl9b A2A3-overexpressing pericytes formed significantly more secondary processes than controls, although their longitudinal length and maximum secondary process length remained unchanged (Supp. Fig. 4I–M). Nevertheless, both cell volume and surface area were significantly increased (Supp. Fig. 4N, O). Thus, reducing either RhoA or myosin II activity promotes a more extensively branched and enlarged pericyte morphology.

Elevated RhoA activity produced a distinct morphological response. RhoA CA-overexpressing pericytes (Fig. 3D–F) progressively elongated along their longitudinal axis between 3 and 14 dpf and are significantly longer than control at 5 and 14 dpf (Fig. 3G). In parallel, their secondary processes shortened more extensively than those of controls, with maximum secondary process length significantly reduced at 5 and 14 dpf (Fig. 3H). The total number of secondary processes remained unchanged (Fig. 3I) whereas their tortuosity is significantly reduced at 14 dpf (Fig. 3J), indicating that elevated RhoA activity promotes secondary process shortening and straightening. However, the accompanying longitudinal elongation offset this loss of secondary process length. Consequently, rather than undergoing uniform compaction, RhoA CA-expressing pericytes tend to have a greater cell volume and exhibit a significantly larger surface area than controls at 14 dpf (Fig. 3K, L).

To exclude the possibility that RhoA-dependent changes in F-actin organisation biased Lifeact-based morphometrics, we repeated key measurements at 5 dpf using membrane-localised mStayGold-CAAX. Corresponding measurements of longitudinal length, maximum secondary process length and secondary process number obtained with the two labels were 97–99% similar across control, RhoA DN- and RhoA CA-expressing vSMCs and pericytes (Supplementary Fig. 5A–L), confirming that Lifeact reliably reports changes in mural cell shape and process induced by RhoA manipulation.

Collectively, our results show that balanced RhoA–actomyosin activity shapes pericyte morphology. Reduced RhoA or myosin II activity maintains an enlarged, highly branched morphology, whereas elevated RhoA activity shortens and straightens secondary processes while promoting longitudinal elongation.

### Retraction of pericyte secondary processes is actomyosin-dependent

We next examined the dynamics of pericyte secondary process remodelling. Closer inspection revealed two classes of pericytes: Type 1 pericytes that display only rod-like, actin-rich bundles (Fig. 4A) and Type 2 pericytes that exhibit a combination of rod-like and lamellar actin structures (Fig. 4B). At 3 dpf, approximately 60% of pericytes were Type 1 and 40% were Type 2; by 5 dpf, Type 2 cells declined to approximately 10% (Fig. 4C). Time-lapse imaging from 80 to 90 hpf showed approximately 70% retraction of lamellar processes, leaving predominantly rod-like structures (Fig. 4D–F, Video 1). Thus, lamellar processes are transient and retract during pericyte maturation.

We subsequently investigated how RhoA activity influences the temporal dynamics of secondary process retraction. RhoA DN-expressing pericytes had fewer Type 1 cells than controls at 3 dpf (Fig. 4C). Although the Type 1 population increased by 5 dpf, this increase was smaller than in controls (Δ = 14% versus 25%), resulting in higher proportion of Type 2 cells. Consistently, lamellar process length changed minimally between 80 and 90 hpf (Fig. 4D, F, G; Video 2). Reduced RhoA activity therefore stabilises lamellar processes and delays acquisition of the Type 1 morphology. RhoA CA-expressing pericytes initially contained a similar proportion of Type 1 cells to controls but also developed a distinct Type 3 morphology, characterised by sparse, short spike-like processes (Fig. 4C, H). Type 3 cells progressively increased and predominated by 5 dpf, with corresponding reductions in Type 1 and Type 2 cells. Time-lapse imaging showed earlier lamellar-process shortening (Fig. 4D, I; Video 3). Because Type 3 cells occurred only following RhoA CA expression, they represent an aberrant morphology associated with elevated contractility.

Thus, actomyosin contractility promotes Type 2-to-Type 1 pericyte transition. Excessive activity produces aberrant Type 3 pericytes, demonstrating that secondary process remodelling requires balanced contractility.

### vSMC contractility stabilises dorsal aorta diameter, whereas pericyte contractility is dispensable for ISV remodelling

As mural cells regulate vessel diameter^13,18,38^, we examined how their contractility and morphology contribute to developmental vascular remodelling^11^. To address this, we generated the following stable transgenic lines: *Tg(6xUAS:lifeact-mCherry)^rk^*^33^ (hereafter referred to as control), *Tg(6xUAS:RhoA T19N-P2A-lifeact-mCherry)^rk^*^34^ (hereafter *Tg(RhoA DN-OE*)), and *Tg(6xUAS:RhoA G14V-P2A-lifeact-mCherry)^rk35^* (hereafter *Tg(RhoA CA-OE)*). These lines were crossed with *TgBAC(pdgfrb:GAL4FF)^ncv24^*zebrafish to drive mural cell–specific transgene expression and with *Tg(kdrl:EGFP)*^s843^ zebrafish to visualise the DA (Fig. 5A–C), aISVs and vISVs (Supp. Fig. 6A–I) at 3, 5 and 14 dpf.

**Figure 5.**
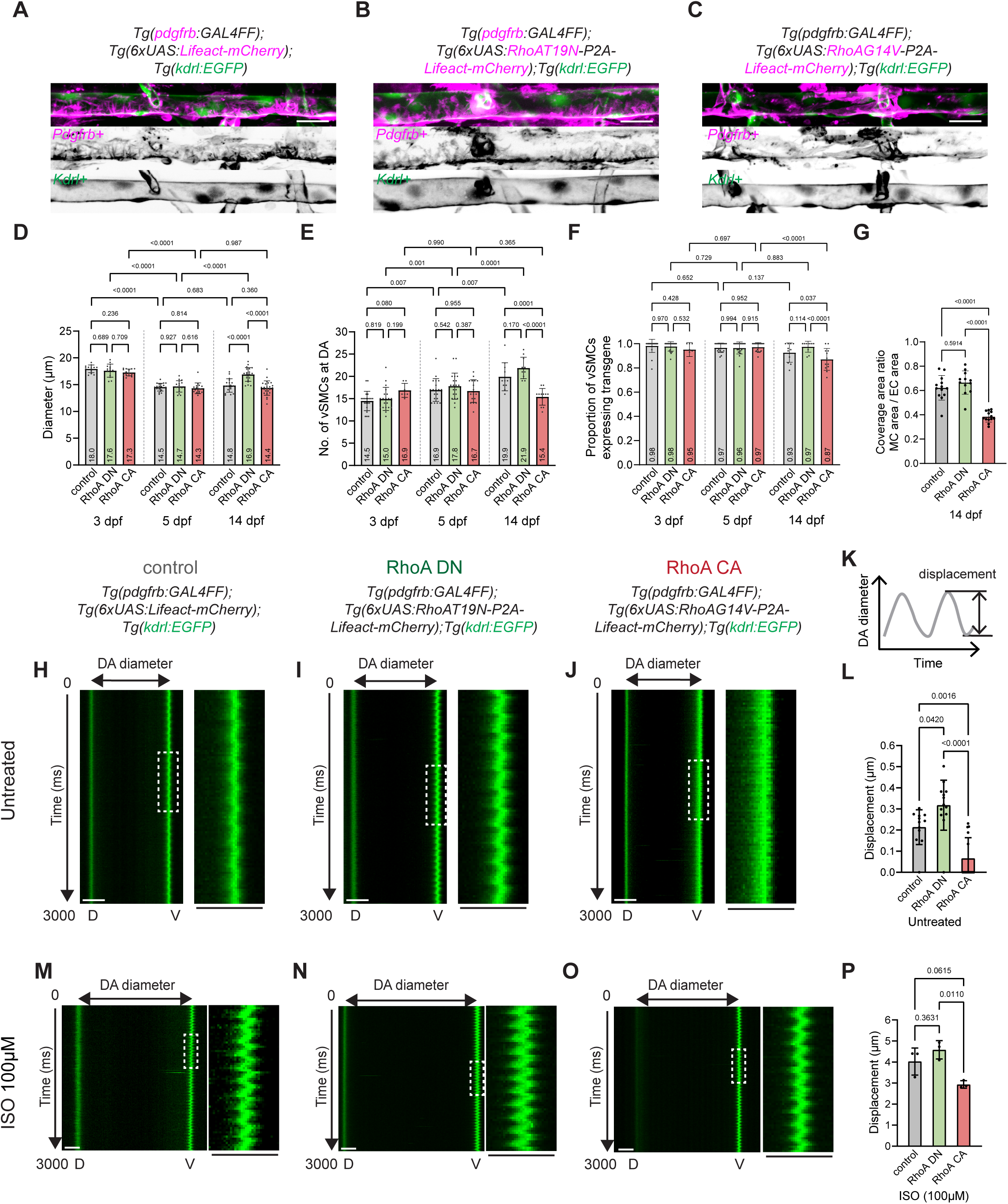
vSMC contractility stabilises dorsal aorta diameter and restricts vessel wall deformation. **A – C** Maximum intensity projection of confocal z-stacks of DA in *TgBAC(pdgfrb:GAL4FF); Tg(6×UAS:Lifeact–mCherry)* (**A**), *TgBAC(pdgfrb:GAL4FF);Tg(6×UAS:RhoA T19N–P2A–Lifeact–mCherry)* (**B**) and *TgBAC(pdgfrb:GAL4FF); Tg(6×UAS:RhoA G14V–P2A–Lifeact–mCherry)* (**C**) embryos at 14 dpf in *Tg(kdrl:EGFP)* background. Magenta, mural cells; green, endothelium. Scale bar, 20 µm. **D** Quantification of DA diameter in control, *Tg(RhoA DN-OE)* and *Tg(RhoA CA-OE)* embryos at 3, 5 and 14 dpf (control: n= 13/20/18 DAs from 13/20/18 embryos at 3/5/14 dpf from 2/3/2 independent experiments; RhoA DN: n= 13/17/21 DAs from 13/17/21 embryos at 3/5/14 dpf from 2/2/2 independent experiments; RhoA CA: n= 14/16/24 DAs from 14/16/24 embryos at 3/5/14 dpf from 2/2/2 independent experiments). Data were analysed by two-way ANOVA with Tukey’s multiple comparisons test. **E** Quantification of vSMC number along a defined DA segment in control, *Tg(RhoA DN-OE)* and *Tg(RhoA CA-OE)* embryos at 3, 5 and 14 dpf. **F** Proportion of vSMCs expressing transgene relative to total vSMCs within a defined segment of the DA in control, *Tg(RhoA DN-OE)* and *Tg(RhoA CA-OE)* embryos at 3, 5 and 14 dpf. **E** control: n= 21/19/11 DAs from 21/19/11 embryos at 3/5/14 dpf; RhoA DN: n= 20/20/10 DAs from 20/20/10 embryos at 3/5/14 dpf; RhoA CA: n= 7/17/12 DAs from 7/17/12 embryos at 3/5/14 dpf from 2 independent experiments. **F** control: n= 8/19/11 DAs from 8/19/11 embryos at 3/5/14 dpf; RhoA DN: n= 8/20/10 DAs from 8/20/10 embryos at 3/5/14 dpf; RhoA CA: n= 7/17/12 DAs from 7/17/12 embryos at 3/5/14 dpf from 2 independent experiments. Data were analysed by two-way ANOVA with Tukey’s multiple comparisons test. **G** Area ratio of endothelial surface within a defined segment of the DA covered by vSMCs in control, *Tg(RhoA DN-OE)* and *Tg(RhoA CA-OE)* embryos at 14 dpf (control: n = 13 DAs from 13 embryos; RhoA DN: n = 12 DAs from 12 embryos; RhoA CA: n = 13 DAs from 13 embryos from 2 independent experiments.) Data were analyzed by ordinary one-way ANOVA with Tukey’s multiple comparisons test. **H – J** Representative kymograph of DA diameter changes in control (**H**), *Tg(RhoA DN-OE)* (**I**) and *Tg(RhoA CA-OE)* (**J**) embryos at 84 - 85 hpf. **K – M** Representative kymographs of DA diameter changes in control (**K**), *Tg(RhoA DN-OE)* (**L**) and *Tg(RhoA CA-OE)* (**M**) embryos after 3-hour treatment with 100 µM Isoprenaline. Scale bar, 5 µm. **N** Corresponding plots of DA diameter in control embryos over time. **O** Average ventral DA wall displacement within 3 seconds at 84–85 hpf (control: n = 12 DAs; RhoA DN: n = 13 DAs; RhoA CA: n = 15 DAs from 2 independent experiments.) **P** Quantification of average ventral DA wall displacement within 3 seconds after 3**-**hour treatment of 100 µM Isoprenaline. (control: n = 5 DAs; RhoA DN: n = 5 DAs; RhoA CA: n = 4 DAs from 2 independent experiments.) Data were analyzed by ordinary one-way ANOVA with Tukey’s multiple comparisons test.

**Figure 6.**
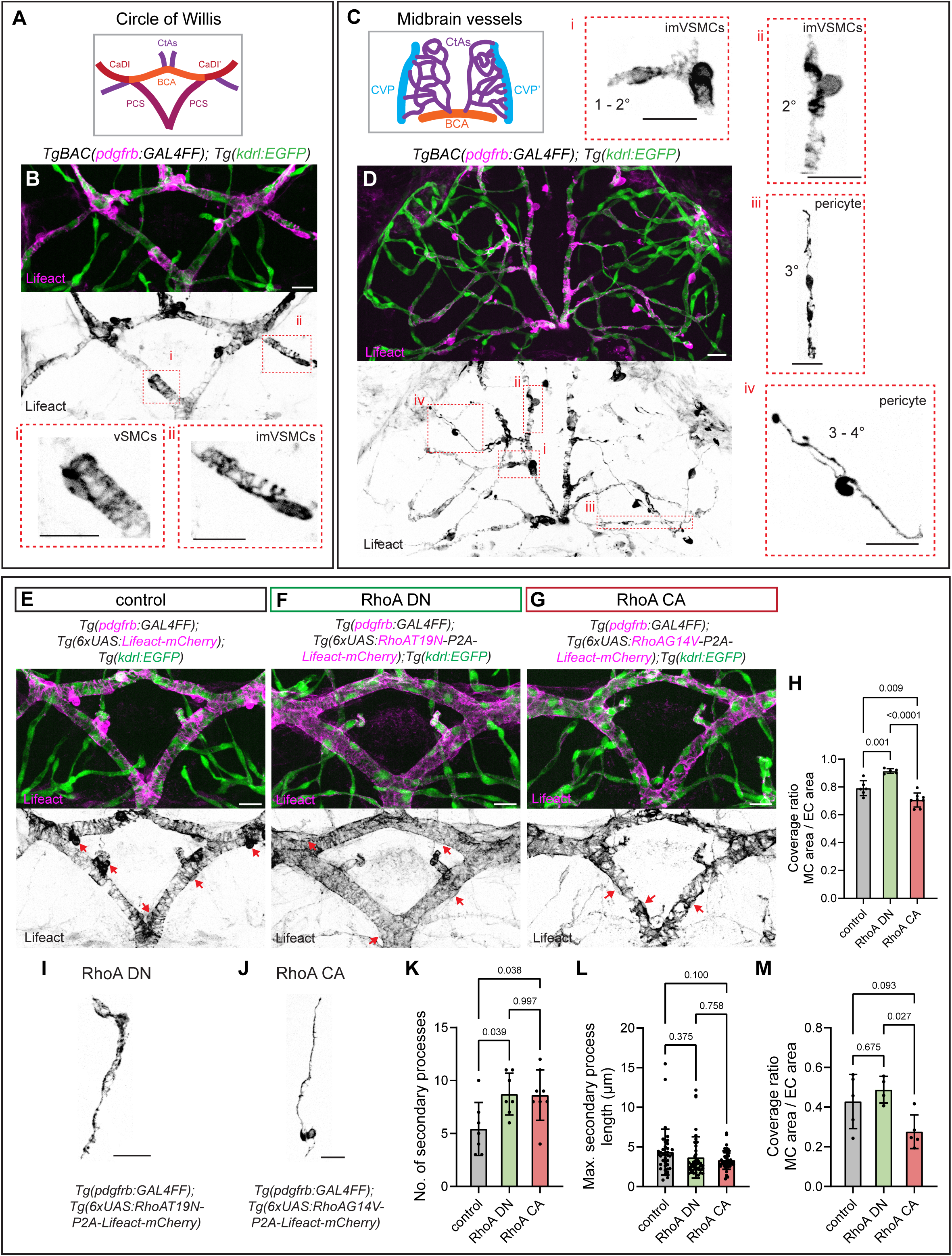
Actomyosin contractility regulates mural cell morphology and vessel coverage in the brain vasculature. **A** Schematic of the CoW and associated vessels. **B** Maximum intensity projection of confocal z-stacks showing vSMCs mosaically expressing Lifeact-mCherry along the CoW at 14 dpf in tissue-cleared *TgBAC(pdgfrb:GAL4FF); Tg(kdrl:EGFP)* embryos. Insets show higher magnification views of vSMC (**i**) and imVSMC (**ii**). **C** Schematic of midbrain vessels showing different branch orders of CtAs. **D** Maximum intensity projection of confocal z-stacks showing mural cell distribution across branch orders of CtAs in tissue-cleared *TgBAC(pdgfrb:GAL4FF); Tg(kdrl:EGFP)* embryos. Insets show higher magnification views of imVSMC along lower order of CtAs **(i, ii**) and pericytes along the higher order CtAs (**iii, iv**). **E – G** Coverage patterns of vSMCs over CoW vessels in *TgBAC(pdgfrb:GAL4FF); Tg(6×UAS:Lifeact–mCherry)* (**E**), *TgBAC(pdgfrb:GAL4FF);Tg(6×UAS:RhoA T19N–P2A–Lifeact–mCherry)* (**F**) and *TgBAC(pdgfrb:GAL4FF); Tg(6×UAS:RhoA G14V–P2A–Lifeact–mCherry)* (**G**) embryos at 14 dpf in *Tg(kdrl:EGFP)* background. **H** Quantification of vSMC coverage of CoW vessels at 14 dpf. Control: n = 6 embryos; RhoA DN: n = 5 embryos; RhoA CA: n = 8 embryos from 2 independent experiment. Data were analyzed by ordinary one-way ANOVA with Tukey’s multiple comparisons test. **I – J** Representative morphology of individual pericytes expressing RhoA DN (**I**) or RhoA CA (**J**). **K - M** RhoA activity modulates pericyte morphology. Quantification of secondary process number (**K**), maximum secondary process length (**L**) and pericyte coverage of CtAs in 14 dpf control, *Tg(RhoA DN-OE)* and *Tg(RhoA CA-OE)* embryos. Control: n = 5 pericytes from 3 embryos, RhoA DN: n = 4 pericytes from 3 embryos, RhoA CA: n = 5 pericytes from 4 embryos from 2 independent experiments. Data were analyzed by ordinary one-way ANOVA with Tukey’s multiple comparisons test. Scale bar, 20 µm. **CoW** Circle of Willis, **CtAs** central arteries, **BCA** basal communicating artery, **CaDI** caudal division of the internal carotid artery, **PCS** posterior communicating segment, **CVP** choroidal vascular plexus.

In control zebrafish, the diameters of the DA, aISVs and vISVs decreased between 3 and 5 dpf and remained stable thereafter (Fig. 5D and Supp. Fig. 6J, K). In *Tg(RhoA DN-OE)* zebrafish, the DA diameter similarly decreased between 3 and 5 dpf but failed to remain constricted thereafter, exhibiting significant dilation at 14 dpf compared with both control and *Tg(RhoA CA-OE)* zebrafish (Fig. 5D). In contrast, both the constriction phase (3–5 dpf) and the subsequent stabilisation phase (5–14 dpf) were preserved in *Tg(RhoA CA-OE)* zebrafish. These findings suggest that vSMC contractility is dispensable for the initial developmental constriction of the DA but is required to maintain its narrowed diameter thereafter, with decreased contractility leading to DA dilation. Notably, the diameters of aISVs and vISVs remained unchanged across all stages examined in *Tg(RhoA DN-OE)* and *Tg(RhoA CA-OE)* zebrafish (Supp. Fig. 6J, K), suggesting that pericyte contractility is dispensable for developmental remodelling of these vessels.

To determine whether the changes in DA diameter could be explained by differences in vSMC abundance or transgene penetrance, we used the *TgBAC(pdgfrb:EGFP)^ncv22^* reporter to quantify the number of EGFP^+^ vSMCs (Fig. 5E) and the proportion of these cells expressing the Lifeact-labelled transgene (Fig. 5F) within a defined DA segment. In *Tg(RhoA DN-OE)* zebrafish, vSMC number and transgene penetrance were indistinguishable from control zebrafish at 3, 5 and 14 dpf. vSMC coverage of the DA surface was also comparable between the two groups at 14 dpf (Fig. 5G). Thus, the DA dilation caused by RhoA DN overexpression cannot be attributed to reduced vSMC abundance, transgene penetrance or vessel coverage, supporting a contribution of reduced vSMC contractility to DA dilation. In *Tg(RhoA CA-OE)* zebrafish, vSMC number and transgene penetrance were comparable to controls at 3 and 5 dpf but were significantly reduced by 14 dpf (Fig. 5E, F), accompanied by decreased vSMC coverage of the DA surface (Fig. 5G). This finding suggests that reduced vSMC coverage may influence the effect of RhoA CA expression on DA diameter, potentially contributing to the control-like DA diameter observed at 14 dpf (Fig. 5D).

We further asked whether the absence of diameter changes in aISVs and vISVs could be explained by altered pericyte abundance or transgene penetrance by using *TgBAC(pdgfrb:EGFP)^ncv22^*reporter (Supp. Fig. 7A – I). Neither parameter differed between *Tg(RhoA DN-OE)* and control zebrafish at 3, 5, or 14 dpf (Supp. Fig. 7J – M). In *Tg(RhoA CA-OE)* zebrafish, pericyte number at vISVs and transgene penetrance in aISVs and vISVs were reduced at 14 dpf, but not at 3 and 5 dpf, when these vessels undergo developmental constriction (Supp. Fig. 7J–M). Despite these later reductions, aISV and vISV diameters of *Tg(RhoA CA-OE)* zebrafish remained comparable to controls at 14 dpf. Developmental remodelling of ISVs therefore proceeds largely independently of pericyte abundance and contractility.

**Figure 7.**
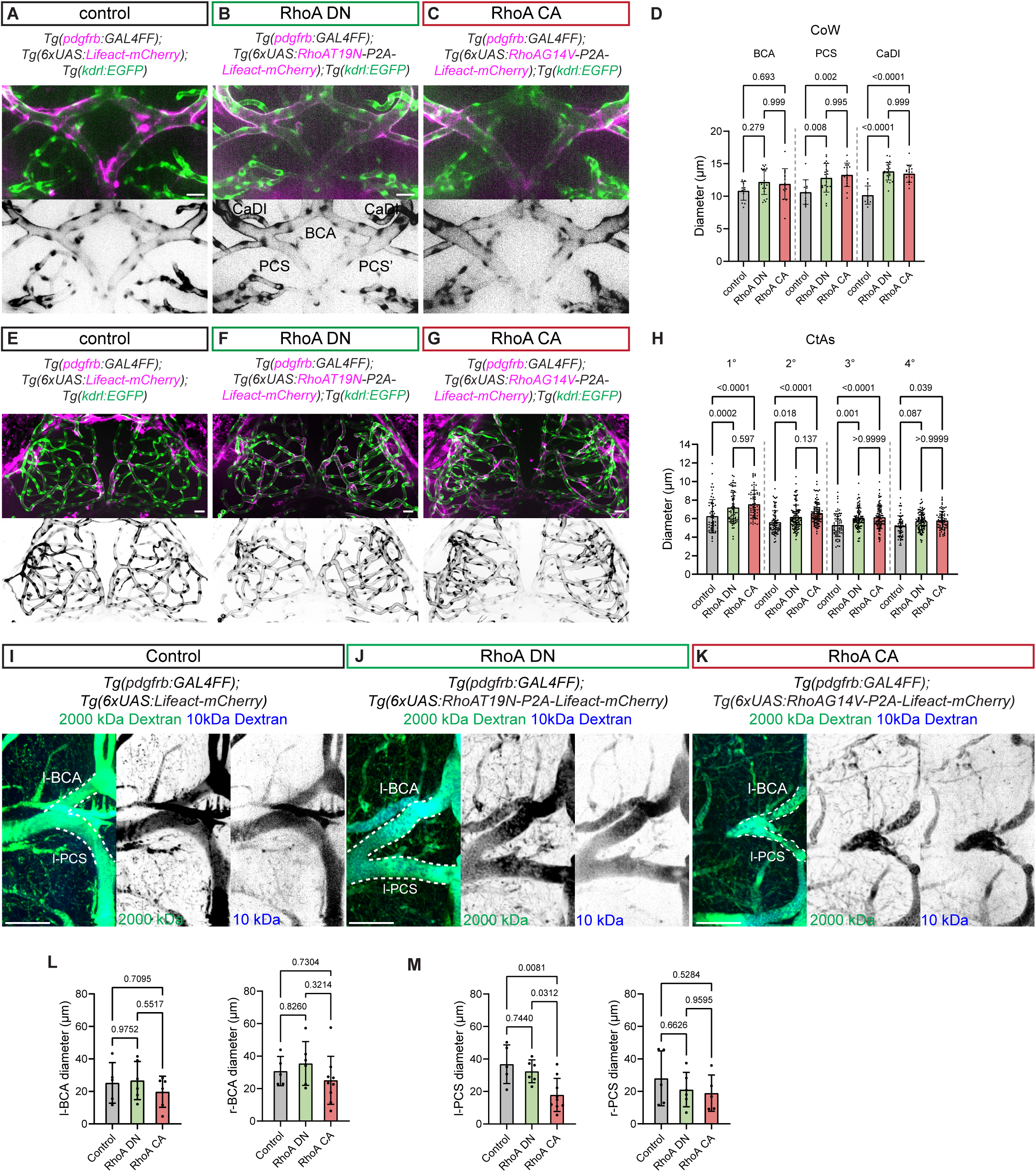
Perturbed mural cell contractility alters brain vessel diameter. **A – C** Maximum intensity projection of confocal z-stacks of CoW at 14 dpf in *TgBAC(pdgfrb:GAL4FF)*; *Tg(6×UAS:Lifeact–mCherry)* (**A**), *TgBAC(pdgfrb:GAL4FF)*; *Tg(6×UAS:RhoA T19N–P2A–Lifeact–mCherry)* (**B**) and *TgBAC(pdgfrb:GAL4FF)*; *Tg(6×UAS:RhoA G14V–P2A–Lifeact–mCherry)* (**C**) embryos in *Tg(kdrl:EGFP)* background. **D** Quantification of vessel diameters in CoW segments (control: n = 12/12/12 BCA/PCS/CaDI from 6 embryos, RhoA DN; n = 16/16/16 BCA/PCS/CaDI from 8 embryos, RhoA CA: n = 18/18/18 BCA/PCS/CaDI from 9 embryos). **E – G** Maximum intensity projection of confocal z-stacks of midbrain vessels at 14 dpf under control (**E**), RhoA DN (**F**) and RhoA CA (**G**) conditions. **H** Quantification of CtA diameter across branch orders (control: n = 32/47/73/61 1°/2°/3°/4° CtAs from 8 embryos, RhoA DN: n = 51/80/56/61 1°/2°/3°/4° CtAs from 8 embryos, RhoA CA: n = 60/82/86/65 1°/2°/3°/4° CtAs from 8 embryos). CoW and CtA diameters were collected from 2 independent experiments and analysed by two-way ANOVA with Tukey’s multiple comparisons test. **CoW** Circle of Willis, **CtAs** central arteries, **BCA** basal communicating artery, **CaDI** caudal division of the internal carotid artery, **PCS** posterior communicating segment. Scale bar, 20 µm. **I – K** Fluorescent tracer assays in 10-month-old *TgBAC(pdgfrb:GAL4FF)*; *Tg(6×UAS:Lifeact–mCherry)* zebrafish (**I**) and 13-month-old *TgBAC(pdgfrb:GAL4FF)*; *Tg(6×UAS:RhoA T19N–P2A–Lifeact–mCherry)* (**J**) and *TgBAC(pdgfrb:GAL4FF)*; *Tg(6×UAS:RhoA G14V–P2A–Lifeact–mCherry)* (**K**) zebrafish. 10 kDa Dextran–Cascade Blue (blue) and 2000 kDa Dextran-Fluorescein (green) were retro-orbitally co-injected. Scale bar, 100 μm. **L – M** Quantification of vessel diameters in CoW segments (control: n = 5/5/5/5 l-BCA/r-BCA/l-PCS/r-PCS from 5 adult fish, RhoA DN; n = 6/6/6/6 l-BCA/r-BCA/l-PCS/r-PCS from 6 adult fish, RhoA CA: n = 6/5/9/9 l-BCA/r-BCA/l-PCS/r-PCS from 10 adult fish). Data were analysed by ordinary one-way ANOVA with Tukey’s multiple comparisons test.

Because *pdgfrb*-driven expression could vary among mural cell populations and developmental stages, we quantified EGFP intensity in *TgBAC(pdgfrb:EGFP)^ncv22^* zebrafish^20^. Reporter intensity increased between 3 and 14 dpf in DA-associated vSMCs and in aISV- and vISV-associated pericytes (Supp. Fig. 7N–P); comparisons were therefore performed within matched vessel beds and developmental stages. We next examined whether the RhoA DN and RhoA CA transgenes were expressed at comparable levels. As each RhoA variant and Lifeact–mCherry are encoded by the same bicistronic construct, Lifeact fluorescence was used as a proxy for transgene expression. At 5 dpf, Lifeact fluorescence was comparable between RhoA DN- and RhoA CA-expressing mural cells associated with the DA, aISVs and vISVs, although both were lower than controls (Supp. Fig. 7Q). Thus, the distinct RhoA DN and RhoA CA vascular phenotypes are unlikely to reflect unequal transgene expression between these two transgenic lines.

Collectively, these results reveal distinct requirements for mural cells during vascular remodelling in the zebrafish trunk. The initial developmental constriction of the DA occurs independently of vSMC contractility, whereas subsequent maintenance of its narrowed diameter requires appropriately regulated vSMC contractile activity. By contrast, the developmental constriction and stabilisation of aISVs and vISVs proceed independently of pericyte contractility and its associated effects on cell morphology.

### vSMC contractility limits DA wall deformation in response to pulsatile blood flow

The DA undergoes cyclic deformation in response to pulsatile blood flow and pressure changes. At 3 dpf, rapid rhythmic changes are readily detected at the ventral DA wall as the vessel expands during systole and recoils during diastole (Fig. 5H; Videos 4–6). To assess the contribution of vSMC contractility, we quantified ventral DA wall displacement at 84 – 85 hpf in control, *Tg(RhoA DN-OE)* and *Tg(RhoA CA-OE)* zebrafish (Fig. 5H–J). Peak-to-trough displacement measured by kymograph analysis (Fig. 5K) was increased in *Tg(RhoA DN-OE)* zebrafish and reduced in *Tg(RhoA CA-OE)* zebrafish, indicating greater and lower wall deformability, respectively (Fig. 5L). We next treated zebrafish with 100 μm isoprenaline hydrochloride (ISO) to examine DA deformation under increased haemodynamic stimulation (Fig. 5M–O). ISO increased DA wall displacement in all three groups. However, displacement was significantly lower in *Tg(RhoA CA-OE)* zebrafish than in control and *Tg(RhoA DN-OE)* zebrafish (Fig. 5P), indicating that elevated vSMC contractility limits circumferential expansion even under increased haemodynamic load.

These results demonstrate that the contractile state of vSMC regulates the mechanical response of the developing DA to pulsatile blood flow: elevated contractility restricts vessel expansion, whereas reduced contractility increases vessel deformability.

### vSMC and pericyte contractility and shape determine mural cell coverage of cerebral vessels

We next extended our analysis to mural cells covering the Circle of Willis (CoW) and central arteries (CtAs) in the zebrafish midbrain. The CoW comprises the basal communicating artery (BCA), the caudal division of the internal carotid artery (CaDI), and the posterior communicating segment (PCS; Fig. 6A)^39,40^. Mosaic labelling reveals that, at 14 dpf, mural cells covering the CoW (Fig. 6B) display bulging cell bodies with actin-rich secondary processes that extend circumferentially around the vessel (Fig. 6B i). Their high *tagln* (Supp. Fig. 8A) and *acta2* expression^24,39^ identifies them as vSMCs. CtAs branch from the BCA and CaDI and further ramify into higher-order vessels (Fig. 6C). Lower-order CtA (1°–2°) mural cells resemble CoW-associated cells but have more prominent cell bodies and discontinuous secondary processes (Fig. 6B ii, D i–ii). Their high *pdgfrb* and low *tagln* expression, together with their intermediate morphology, identify them as transitional or intermediate vSMCs (imSMCs). In contrast, higher-order CtA (≥3°) mural cells lack *tagln* (Supp. Fig. 8B) and *acta2* but express the pericyte markers, *ndufa4l2a* and *abcc9*^24,33,33^. Morphologically, they are characterised by protruding cell bodies, elongated primary processes that extend longitudinally along the vessel and few secondary processes spanning across the vessel circumference (Fig. 6D iii–iv), resembling thin-strand pericytes present in mouse brain capillaries^8,41^.

**Figure 8.**
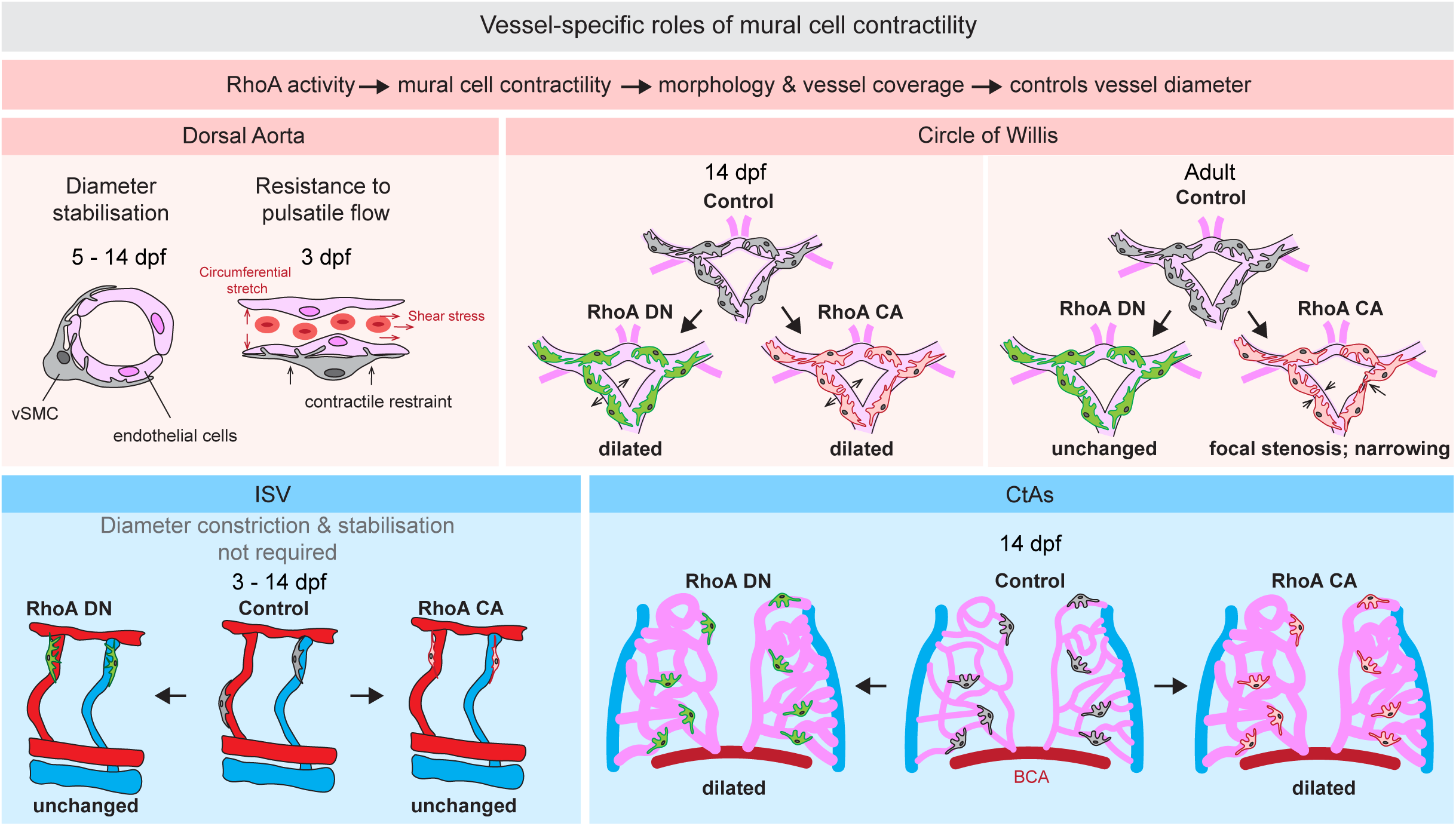
Model illustrating how cell contractility regulates mural cell morphology and vascular diameter during zebrafish development. During development, DA-associated vSMCs and ISV-associated pericytes undergo morphological remodelling characterised by reductions in cellular process length and number and overall cell size. RhoA-dependent actomyosin contractility regulates these morphological changes and their functional consequences in a vessel-specific manner. In the DA, vSMC contractility is dispensable for the initial developmental constriction between 3 and 5 dpf, but is required for subsequent DA diameter stabilisation between 5 and 14 dpf. vSMC contractility also limits DA wall deformation in response to pulsatile blood flow, thereby contributing to vascular tone and mechanical stability. In the cerebral vasculature, vSMC and pericyte contractility contributes to diameter regulation of both the CoW and CtAs, respectively. Perturbation of RhoA activity alters mural cell morphology and vessel coverage and consequently disrupts normal cerebral vessel calibre. Together, these findings support a model in which RhoA-dependent mural cell contractility regulates vascular diameter not through contractile force alone, but through coordinated effects on cell morphology, vessel coverage and mechanical constraint in a vessel- and cell type-specific manner.

We next investigated how contractility influences vSMC morphology and coverage of CoW and CtAs. At 14 dpf, vSMCs of control zebrafish exhibit protrusive actin-rich cell bodies and unevenly distributed circumferential processes, leaving approximately 20% of the vessel surface uncovered (Fig. 6E and H). RhoA DN-overexpression flattened the cell body and promoted secondary processes spreading, significantly increasing vessel coverage in *Tg(RhoA DN-OE)* zebrafish (Fig. 6F and H). Conversely, RhoA CA-overexpression produced disorganised secondary processes and more discontinuous and fragmented coverage, significantly reducing coverage relative to both control and *Tg(RhoA DN-OE)* zebrafish (Fig. 6G, H).

To specifically assess pericyte morphology, we restricted morphometric analysis to thin-strand pericytes associated with ≥3° branches. Both RhoA DN- and RhoA CA-expressing pericytes formed more secondary processes than controls, without changes in maximum secondary-process length (Fig. 6I–L). However, RhoA CA-expressing cells had narrower primary processes and a simpler morphology, resulting in significantly lower vessel coverage than RhoA DN-expressing cells (Fig. 6M).

These findings demonstrate that actomyosin contractility is also a key regulator of brain vSMC and pericyte morphology and vessel coverage in zebrafish brain.

### Perturbation of mural cell contractility alters cerebral vessel diameter in developing and adult zebrafish

To determine whether changes in mural cell contractility, shape and vessel coverage influence the diameter of CoW and CtAs, we analysed live zebrafish at 14 dpf (Fig. 7A-H). In the CoW, control zebrafish exhibit relatively uniform vessel diameters across BCA, PCS and CaDI (Fig. 7A, D). Strikingly, both reduced vSMC contractility in *Tg(RhoA DN-OE)* zebrafish and enhanced vSMC contractility in *Tg(RhoA CA-OE)* zebrafish resulted in increased vessel diameters across PCS and CaDI (Fig. 7B-D). A similar trend was observed in the CtA (Fig. 7E–H). In control zebrafish, CtA diameter progressively decreased from proximal (1°) to distal (4°) branches, reflecting the hierarchical organisation of the vascular tree (Fig. 7H). However, the overexpression of both RhoA DN and RhoA CA led to significant increases in vessel diameter across multiple branch orders, particularly in proximal (1°–3°) branches (Fig. 7F-H). Notably, *Tg(RhoA CA-OE)* zebrafish displayed a shift towards higher-order branches in the midbrain, indicating an increased vascular branching complexity compared to controls (Supp. Fig. 9A). This suggests that enhanced contractility is associated not only with changes in brain vessel diameter but also with vascular patterning.

We next examined whether cerebral vessel calibre changes persisted into adulthood. The left and right BCA and right PCS showed no significant differences among control, *Tg(RhoA DN-OE)* and *Tg(RhoA CA-OE)* zebrafish, whereas the left PCS was significantly narrower in adult *Tg(RhoA CA-OE)* zebrafish (Fig. 7I - M). Focal CoW stenoses were also detected in 7 out of 10 *Tg(RhoA CA-OE)* brains (Supp. Fig. 10A–C). Thus, the widespread cerebral vessel dilation in *Tg(RhoA CA-OE)* larvae was replaced by regionally restricted narrowing in adults.

Because mural cells contribute to blood-brain-barrier integrity, we further examined whether sustained alterations in mural cell contractility compromised vascular permeability in adult zebrafish. Vascular permeability was assessed by co-injecting 2000-kDa and 10-kDa dextran tracers. Normalised extracellular 10-kDa dextran fluorescence did not differ among control, *Tg(RhoA DN-OE)* and *Tg(RhoA CA-OE)* zebrafish (Supp. Fig. 10D), indicating no detectable vascular leakage following reduced nor elevated mural cell RhoA activity.

These findings show that mural cell regulation of cerebral vessel calibre depends on vascular location and developmental stage (Fig. 8). In larvae, both reduced and elevated RhoA activity caused dilation, consistent with a balance between mural cell contractility and vessel coverage: reduced contractility weakens vessel restraint, whereas RhoA CA-induced cell compaction reduces the area over which restraint is exerted. In adults, widespread dilation was replaced by focal CoW narrowing and stenosis following elevated RhoA activity, without detectable vascular leakage, suggesting that the relationship between mural cell contractility and cerebral vessel calibre changes with vascular maturation.

## Discussion

Mural cells are widely regarded as contractile regulators of vessel diameter. Our findings support a broader model in which vessel calibre reflects the integration of mural cell contractility, morphology and coverage. By remodelling how mural cells engage the vessel wall, contractile activity can alter vessel diameter without producing a simple linear relationship between force generation and constriction.

Actomyosin contractility drives developmental changes in trunk mural cell shape. vSMCs and pericytes initially form extensive actin-rich processes that progressively retract, reducing cell size and converting vSMCs to a compact morphology while shifting pericytes from Type 2 to Type 1 states (Figs. 1 and 4A–C). Reducing RhoA or myosin II activity preserved larger, more branched cells, whereas elevated RhoA activity accelerated vSMC compaction and selectively remodelled pericyte processes. A related transition occurs in the postnatal mouse retina, where stellate, proliferative pericytes become quiescent and elongated as PI3K signalling declines^23^. The loss of lamellar processes in zebrafish may therefore reflect a conserved maturation programme involving cytoskeletal signalling, although the relationship between PI3K and RhoA was not tested here.

Our findings distinguish the vSMC-independent initial narrowing of the DA from the vSMC-dependent maintenance of its constricted diameter. Altering RhoA activity did not prevent the initial reduction in DA diameter between 3 and 5 dpf (Fig. 5 A-D), consistent with previous evidence that endothelial cell shape changes^12^ and basement-membrane assembly^6^ contribute to early vessel narrowing. After this phase, reduced RhoA activity led to DA dilation, whereas elevated RhoA activity maintained the narrowed diameter without inducing further constriction between 5 and 14 dpf. Reduced RhoA activity also increased deformation of the DA wall during pulsatile blood flow, while elevated RhoA activity reduced deformation, supporting a role for vSMCs in stabilising vessel diameter and resisting circumferential stretch (Fig. 5H-P). Interpretation of the RhoA CA phenotype is complicated by the accompanying reduction in vSMC abundance and vessel coverage at 14 dpf. Its effect on DA diameter therefore cannot be attributed to increased intrinsic contractility alone, because RhoA activity, cell number and vessel coverage change concurrently in this model.

The cerebral vasculature showed a different relationship between mural cell contractility and vessel diameter. At 14 dpf, both increased and decreased RhoA activity dilated the BCA, PCS and CaDI (Fig. 7A-H). Reduced activity increased vSMC spreading and coverage but weakened contractile restraint, whereas elevated activity compacted vSMCs and reduced the surface over which force could be transmitted (Fig. 6F-H). Effective vessel constraint therefore appears to require an appropriate balance between cellular force and coverage. These larval phenotypes were not broadly maintained in adults: most CoW segments were comparable to controls, whereas RhoA CA expression was associated with left PCS narrowing and focal CoW stenosis (Fig. 7I-M). This transition from widespread larval dilation to restricted adult narrowing suggests that the effect of mural cell RhoA activity changes as coverage, extracellular matrix and vessel wall architecture mature. Because adult mural cell morphology and coverage were not quantified, the basis of this stage-dependent change remains unresolved.

The contribution of pericytes to capillary diameter remains debated. Pericyte ablation can dilate mouse brain capillaries, and activation has been reported to induce rapid constriction, whereas other studies find no effect of pericyte depolarisation on diameter^13,41,42^. Our experiments instead examine developmental remodelling over days. In zebrafish, both RhoA perturbations dilated CtAs, although dilation of upstream CoW vessels may contribute to this phenotype. By contrast, neither ISV constriction nor stabilisation was affected by altered pericyte contractility or morphology (Supplementary Fig. 6J, K), despite extensive actin-rich processes previously proposed to constrict vessels mechanically^43^. This agrees with normal ISV remodelling after reduced pericyte recruitment in *pdgfrb*-depleted zebrafish^11^ and indicates that endothelial mechanisms predominate in this vascular bed. Pericyte control of calibre is therefore vessel-bed dependent rather than universal as demonstrated by pancreatic islet pericytes, which express contractile markers and regulate capillary flow^16^. Sparse coverage may limit mechanical influence in trunk ISVs, whereas the denser mural organisation of cerebral vessels may permit greater regulation through pericyte morphology and coverage.

The function of the elaborate secondary processes formed by trunk pericytes remains unclear because their remodelling did not alter ISV diameter. Perivascular fibroblasts, which are pericyte precursors^44^, may use these secondary processes to contact and migrate onto ISVs before retracting after attachment. Alternatively, secondary processes may organise the basement membrane deposited between pericytes and endothelial cells during vascular maturation.

Several limitations should be considered. RhoA regulates migration, proliferation, gene expression and intercellular signalling in addition to actomyosin contractility ^45^.Therefore, some of the vascular phenotypes caused by RhoA manipulation may result from non-contractile effects, including altered communication between mural cells and endothelial cells. Although zebrafish data do not support a prominent mural cell-derived Angiopoetin mechanism, we did not examine endothelial Angiopoietin–Tie signalling and cannot exclude its indirect contribution^46–48^. We inferred contractile state from the established activities of RhoA T19N and RhoA G14V and their effects on actin organisation and morphology, but did not directly measure RhoA activity or force, or quantify endothelial cell number and shape. Altered mural cell force and coverage could therefore affect vessel diameter indirectly through endothelial remodelling. The reduced number of *pdgfrb*-labelled vSMCs in RhoA CA-expressing zebrafish at 14 dpf remains unexplained and could reflect altered cell maintenance, survival or *pdgfrb* reporter expression. Moreover, continuous RhoA manipulation in *pdgfrb*-expressing cells cannot distinguish immediate mechanical effects from accumulated developmental changes. Finally, RhoA CA simultaneously increases intrinsic contractility and reduces coverage, preventing their individual contributions to vessel diameter from being resolved. Temporally controlled and independent manipulation of these parameters will be required.

In summary, mural cell contractility regulates vessel diameter through combined effects on force, morphology and coverage. vSMCs stabilise arterial diameter and vessel-wall mechanics, whereas pericyte influence depends on vascular context and the extent of vessel coverage. This framework explains why increasing or decreasing RhoA activity can produce similar dilation in some vessels yet different effects in others, and suggests how dysregulated RhoA-dependent remodelling may contribute to vascular disease, including hypertension^49,50^.

## Methods

### Zebrafish maintenance and stocks

All animal experiments were approved by the Institutional Animal Care and Use Committee at RIKEN Kobe Branch (IACUC). Zebrafish (Danio rerio) were raised and staged according to established protocols^51^. Transgenic lines used in this work are *Tg(kdrl:EGFP)^s843^* ^52^, *Tg(fli1:Lifeact-EGFP)^zf495^* ^53^*, TgBAC(pdgfrb:EGFP)^ncv22^* ^20^*, TgBAC(pdgfrb:GAL4FF)^ncv24^* ^20^ *Tg(fli1:myr-mCherry)^ncv1^* ^54^, *TgBAC(tagln:EGFP)^ncv25^* ^20^, *Tg(6xUAS:lifeact-mCherry)^rk33^* (this study), *Tg(6xUAS:RhoA T19N-P2A-lifeact-mCherry)^rk34^* (this study) and *Tg(6xUAS:RhoA G14V-P2A-lifeact-mCherry)^rk35^* (this study). Zebrafish embryos were collected within 30 minutes of divider removal and were allowed to develop at 28.5 °C to the appropriate stage.

### Plasmid construction

Plasmids were generated using In-Fusion^®^ HD Cloning (Takara). A P2A-containing intermediate, pDestTol2-6xUAS: TagBFP-P2A-Lifeact-mCherry, was first generated from pDestTol2-6xUAS-Lifeact-mCherry and subsequently used to construct pDestTol2-6xUAS RhoA T19N-P2A-Lifeact-mCherry, pDestTol2-6xUAS RhoA G14V-P2A-Lifeact-mCherry and pDestTol2-6xUAS Myl9bA2A3-P2A-Lifeact-mCherry.To generate pDs-6xUAS:mStayGold-CAAX, pDs-6xUAS:aqp8a1-P2A-EGFP-CAAX was PCR-linearised to remove *aqp8a1*-P2A-EGFP and assembled with the mStayGold-CAAX coding sequence. For all cloning steps, backbones and insert fragments were PCR-amplified using CloneAmp^®^ HiFi PCR Premix (Takara). Amplification and linearization were verified by 1% TAE agarose gel electrophoresis. PCR products were either used directly or purified using the QIAquick^®^ Gel Extraction Kit (QIAGEN) when necessary. Details of vectors are provided in Supplementary Data.

### Mosaic expression of constructs and generation of transgenic zebrafish lines

For plasmid overexpression, 2 nl of a mixture containing 50–100 ng plasmid DNA, 100–200 ng Tol2 transposase mRNA, phenol red as a tracer and Milli-Q water to volume was co-injected into one-cell stage zebrafish embryos. Tol2 transposase mRNA was transcribed from NotI-linearized pCS-TP vector (a gift from Koichi Kawakami, National Institute of Genetics, Japan) using the mMESSAGE mMACHINE SP6 kit (Invitrogen). For mosaic expression of transgenes, embryos were analysed at 3, 5 or 14 dpf after injection. To generate *Tg(6xUAS:lifeact-mCherry)^rk33^, Tg(6xUAS:RhoA T19N-P2A-lifeact-mCherry)^rk34^* and *Tg(6xUAS:RhoA G14V-P2A-lifeact-mCherry)^rk35^* lines, injected embryos were raised to adults and screened for founders.

### Image acquisition and processing

For live confocal imaging, embryos were embedded in 0.8% low-melting-point agarose (Bio-Rad) prepared in E3 medium containing 0.16 mg/mL Tricaine (Sigma-Aldrich) and 0.003% phenylthiourea (Sigma-Aldrich) to minimise movement and pigmentation, respectively. Confocal z-stacks were collected on an inverted Olympus IX83/Yokogawa CSU-W1 spinning-disk confocal microscope equipped with a Zyla 4.2 CMOS camera (Andor) and Olympus UPLSAPO 40x/NA 1.25 or 30x/NA 1.05 silicone oil immersion objectives. Bright-field images were obtained using a Leica M205FA microscope. Raw images of single mural cell were deconvoluted with ™Huygens Spinning Disk Deconvolution, following the standard protocol established in the unit. The deconvoluted images were subsequently processed using FIJI software (NIH).

### Mural cell morphometrics analysis

Deconvoluted images were processed using IMARIS (v.9.0.0, BitPlane, Zurich, Switzerland), a 3D image visualization and processing software. Images were binarized according to a built-in, self-adapting algorithm. Triangular surface meshes of vessels and mural cells were created with a 0.50 µm and 0.16 µm surface detail resolution setting, respectively. Exported wrl mesh files were converted and downsampled to compact size .ply files via Meshlab (Ver. 2023.12) and PyMeshLab (2023.12. post3) Python library.

Software architecture and implementation details are presented in the Supplementary Figure 2. Briefly, we developed a Python-based tool that enables interactive selection of start and end points for mural cell processes, including both primary and secondary processes, as described previously^43^. Secondary processes were segmented and analysed by defining start points at junctions where the soma or primary processes meet the secondary processes, and end points at their distal tips. For centerline extraction, the cost function of the fast-marching algorithm was empirically set to 1/R to balance propagation across inter-anastomosing segments of secondary processes. Following centerline extraction and automatic tubular mask–based mesh segmentation, users were able to review and manually correct the results. Bifurcations were calculated using VMTK functions that are based on the local Voronoi diagram diagram^55^. Each segment between successive bifurcations was treated as an individual branch for parameter calculation, with the branch closest to the start point defined as first order. In this study, branches originating from the same start point (within a distance smaller than the centerline point interval) and spatially separated from other groups were classified as a single secondary process. We treated inter-anastomose structures as separate branches that close but not adjacent, to prevent software kernel death and keep secondary process topology. Maximum secondary process length was defined as the longest path from the start points to the most distal end point, determined using a two-pass Dijkstra search^56^. For the number of secondary processes. The total number of secondary processes was defined by the number of start points.

### Tracking of lamellar processes in trunk pericytes

To track and quantify dynamic changes in lamellar processes of pericytes, live *Tg(kdrl:EGFP)^s843^*; *TgBAC(pdgfrb:GAL4FF)^ncv24^*zebrafish were outcrossed with *Tg(6xUAS:lifeact-mCherry)^rk33^, Tg(6xUAS:RhoA T19N-P2A-lifeact-mCherry)^rk34^*or *Tg(6xUAS:RhoA G14V-P2A-lifeact-mCherry)^rk35^*. Embryos were collected and subjected to time-lapse imaging starting at 80 hpf, with a total imaging duration of 10 hours. For each process, a straight line was drawn along the axis of the lamella, starting from the region proximal to the soma and primary process and extending to the distal edge of the lamella. Kymographs were generated along this line to track temporal changes in lamellar process extension and retraction over the imaging period. To quantify changes in lamellar process length, measurements were obtained at the initial time point (0 h) and after 10 hours. The percentage change in length was calculated relative to the initial measurement. Multiple processes per cell and multiple cells per embryo were analysed to ensure robust quantification.

### Quantification of mural cell fluorescence intensity and transgene expression

Fluorescence intensity measurements were performed using Fiji. To quantify the expression level of the *Pdgfrb* reporter, individual mural cells, including pericytes and vSMCs, associated with ISVs no. 12–15 and the intervening segment of the DA were identified. The entire cell perimeter was manually outlined using the Freehand Selection tool in Fiji, and the mean EGFP fluorescence intensity within the selected cellular region was measured. To quantify Lifeact–mCherry fluorescence intensity, the outer contour of the cell nucleus was manually outlined using the Freehand Selection tool, and the mean mCherry fluorescence intensity within the nuclear region was measured.

### Quantification of the proportion of transgene-expressing mural cells

For pericytes, the proportion of transgene-expressing cells was quantified for each ISV from no. 12 to no. 15. The number of mCherry-positive pericytes was divided by the total number of EGFP-positive pericytes on the same ISV, and the ratio was calculated. For vSMCs, the proportion of transgene-expressing cells was quantified within the DA segment corresponding to ISVs no. 12–15. The number of mCherry-positive vSMCs covering the DA surface was divided by the total number of EGFP-positive vSMCs within the same DA segment.

### Quantification of blood vessel diameter

*Tg(kdrl:EGFP)^s843^*; *TgBAC(pdgfrb:GAL4FF)^ncv24^*zebrafish were outcrossed with *Tg(6xUAS:lifeact-mCherry)^rk33^, Tg(6xUAS:RhoA T19N-P2A-lifeact-mCherry)^rk34^* or *Tg(6xUAS:RhoA G14V-P2A-lifeact-mCherry)^rk35^* line. Embryos were collected and imaged at 3, 5 and 14 dpf. Confocal z-stacks were acquired with an Olympus UPLSAPO 40×/NA1.25 silicone oil immersion objective. For DA diameter quantification, 6–8 measurements were obtained between ISVs no. 10 and 15 for each embryo and averaged. For ISV diameters, six measurements were acquired along each vessel (ISVs no. 10–15) and averaged. For BCA, PCS and CaDI diameters, each bilateral vessel was measured separately. For each side, over three measurements were obtained and averaged to yield a single value per vessel. To quantify CtA diameter, midbrain vessels between the BCA and CVP were first hierarchically classified based on branching order. For each interbranch segment, over three measurements were obtained and averaged to determine the diameter of that segment. Measurements were performed using Fiji (NIH).

### Quantification of vSMC coverage area on arteries

*Tg(kdrl:EGFP)^s843^*; *TgBAC(pdgfrb:GAL4FF)^ncv24^*zebrafish were outcrossed with *Tg(6xUAS:lifeact-mCherry)^rk33^, Tg(6xUAS:RhoA T19N-P2A-lifeact-mCherry)^rk34^* or *Tg(6xUAS:RhoA G14V-P2A-lifeact-mCherry)^rk35^* line. Embryos were collected and analysed at 14 dpf. For quantification of the coverage area of vSMCs on vessels within the CoW, imaging was performed on tissue-cleared samples (CUBIC Trial Kit, Fujifilm), enabling improved optical penetration and better visualisation of vSMC morphology. Regions of interest (ROIs) corresponding to the EC area were first manually delineated using the freehand selection tool in Fiji, and the total EC area was measured. Subsequently, the Lifeact–mCherry channel was thresholded to segment mural cell structures within the CoW, thereby generating a binary mask representing vSMC distribution. Segmentation signals outside the predefined ROI were excluded to restrict the analysis to the vessel area. The area occupied by vSMCs within the ROI was then quantified, and the coverage ratio was calculated as the fraction of vSMC-positive area relative to the total EC area. The coverage area within the DA was quantified using a similar approach. However, images were acquired from live embryos, and the analysis was performed as described above.

### Measurement of DA wall displacement

*Tg(kdrl:EGFP)^s843^*; *TgBAC(pdgfrb:GAL4FF)^ncv24^*zebrafish were outcrossed with *Tg(6xUAS:lifeact-mCherry)^rk33^, Tg(6xUAS:RhoA T19N-P2A-lifeact-mCherry)^rk34^* or *Tg(6xUAS:RhoA G14V-P2A-lifeact-mCherry)^rk35^* line. Embryos were imaged at 84 – 85 hpf. To increase haemodynamic forces, embryos were treated with 100 µM isoprenaline hydrochloride (Sigma-Aldrich, I5627-5G) in 12-well plates for 3 hours before imaging, with exposure maintained during live imaging by adding E3 buffer containing 100 µM isoprenaline hydrochloride over the agarose gel. Time-lapse imaging was performed with 10 ms interval for a total duration of 20 s. Images were acquired at a single focal plane positioned at the middle plane of the DA. To quantify movement of the DA wall over time, a line ROI was drawn across the vessel to generate a kymograph in Fiji. The oscillatory movement of the DA wall during cardiac cycles was manually traced on the kymograph. The DA diameter was measured for each time frame to track temporal changes. From these measurements, the maximum displacement of the ventral side of the DA over time was determined and averaged.

### Tracer dye injections and imaging

For all injections at adult stage, 10 kDa Dextran–Cascade Blue (Thermo Fisher Scientific, D1976) was mixed and co-injected with 2,000 kDa Dextran–Fluorescein (Thermo Fisher Scientific, D7137). The 2,000 kDa dextran tracer was used as an internal reference to normalize fluorescence signals and control for variation in the amount of tracer delivered intravenously across imaging experiments. Both tracers were used at a final concentration of 5 mg/ml. Intravenous injections in adult zebrafish were performed retro-orbitally using a glass capillary needle, as previously described^57^. At 2 h post-injection, brains were dissected and fixed overnight in 2% PFA. Whole brains were subsequently cleared using a CUBIC-based tissue-clearing method. Cleared brains were imaged using an Olympus FV3000 confocal microscope equipped with a 10×/0.4 NA objective, with image stacks acquired at a 1 µm z-step interval.

### Tracer intensity

Tracer intensity quantification was performed in Fiji. Tracer accumulation within the brain parenchyma was quantified for 10 kDa dextran. Mean fluorescence intensity was measured from four regions of interest (ROIs) within the midbrain parenchyma using 100-µm-thick maximum-intensity projections (MIPs) at CoW. Parenchymal tracer signals were normalized to the vascular signal of 2,000 kDa dextran, which was quantified as the mean fluorescence intensity from four ROIs positioned at CoW. For each sample, normalized tracer intensity was calculated as the mean parenchymal fluorescence intensity of the 10 kDa tracer divided by the mean luminal fluorescence intensity of the 2,000 kDa dextran tracer.

## Statistical analysis

All statistical analyses and graphical representations were conducted using GraphPad Prism 10 software. Further details on statistics for each data set can be found in the corresponding figure legend. Comparisons among more than two groups for a single factor were performed using one-way ANOVA followed by Tukey’s post hoc test. For analyses involving two factors, two-way ANOVA followed by Tukey’s post hoc test was used. Categorical data were analysed using Fisher’s exact test. Exact P values are reported in the figures.

## Data availability

All data supporting the findings of this study are available within the article and its Supplementary Information files. The custom code used for image processing and morphometric analysis is available in GitHub (https://github.com/HedeleZeng/Mural-VISTA.git).

## Acknowledgements

We thank members of the Phng laboratory for valuable discussions and suggestions; Rajrishi Kumar, Emi Taniguchi and the RIKEN BDR Research Aquarium for technical support; Yan Chen for advice on Fiji macros; and Yasushi Okada for sharing plasmid. This work was supported by intramural funding from RIKEN BDR (to L.-K.P.), JSPS Grants-in-Aid for Scientific Research grants (22H022624 and 22H05168 to L-K.P); RIKEN Junior Research Associate Programme (to M.H.), CSC–OU joint scholarship to M.H.) and Pioneering Research Initiated by the Next Generation (SPRING) by Japan Science and Technology Agency (JPMJSP2108 to H.Z.).

## Author contributions

M.H. and L.-K.P. conceived the study. M.H., R.C. and K.A. performed experiments. H.Z. and Y.T.M. conducted mural cell morphometrics analysis and prepared the corresponding figures and text. M.H. analysed all other data and prepared the figures and movies. M.H. and L.-K.P. wrote the manuscript with input from all authors. L.-K.P. acquired funding and supervised the project. All authors discussed the results and commented on the manuscript.

## Competing Interests

The authors declare no competing interests.

## Notes

### Competing Interest Statement

The authors have declared no competing interest.

### Summary of Updates

We have in the revised manuscript clarified the anatomical, morphological and molecular criteria used to distinguish vascular smooth muscle cells and pericytes; independently validated the principal Lifeact based morphometric measurements using the membrane localised reporter mStayGoldCAAX; introduced a downstream perturbation of nonmuscle myosin II, which reproduced key features of the RhoA dominant negative phenotype and provides additional evidence linking RhoA/myosin II activity to mural-cell shape; added quantitative analyses of secondary process tortuosity, transgene expressing mural-cell coverage, developmental changes in pdgfrb reporter activity and relative transgene expression; extended the study to adult cerebral vessels, revealing regionally restricted narrowing and focal stenosis following elevated RhoA activity, while finding no detectable impairment of blood brain barrier integrity; and expanded the Discussion to address how changes in mural cell force, morphology and vessel coverage may influence vascular calibre, while acknowledging possible contributions from endothelial remodelling and mural cell endothelial signalling, including Angiopoietin/Tie signalling.

## Reference

1. Gaengel, K., Genové, G., Armulik, A. & Betsholtz, C. Endothelial-Mural Cell Signaling in Vascular Development and Angiogenesis. ATVB 29, 630–638 (2009).

2. Armulik, A., Genové, G. & Betsholtz, C. Pericytes: Developmental, Physiological, and Pathological Perspectives, Problems, and Promises. Developmental Cell 21, 193–215 (2011).

3. Ando, K., Ishii, T. & Fukuhara, S. Zebrafish Vascular Mural Cell Biology: Recent Advances, Development, and Functions. Life 11, 1041 (2021).

4. Siekmann, A. F. Biology of vascular mural cells. Development 150, dev200271 (2023).

5. Whitesell, T. R. et al. An α-Smooth Muscle Actin (acta2/αsma) Zebrafish Transgenic Line Marking Vascular Mural Cells and Visceral Smooth Muscle Cells. PLoS ONE 9, e90590 (2014).

6. Stratman, A. N. et al. Mural-Endothelial cell-cell interactions stabilize the developing zebrafish dorsal aorta. Development dev.143131 (2016) doi:10.1242/dev.143131.

7. Sweeney, M. D., Ayyadurai, S. & Zlokovic, B. V. Pericytes of the neurovascular unit: key functions and signaling pathways. Nat Neurosci 19, 771–783 (2016).

8. Grant, R. I. et al. Organizational hierarchy and structural diversity of microvascular pericytes in adult mouse cortex. J Cereb Blood Flow Metab 39, 411–425 (2019).

9. Eilken, H. M. et al. Pericytes regulate VEGF-induced endothelial sprouting through VEGFR1. Nat Commun 8, 1574 (2017).

10. Crawshaw, J. R., Flegg, J. A., Bernabeu, M. O. & Osborne, J. M. Mathematical models of developmental vascular remodelling: A review. PLoS Comput Biol 19, e1011130 (2023).

11. Chen, Y. et al. Circumferential actomyosin bundles anchored by CCM1 drive endothelial cell contraction and vessel constriction. Nat Commun 17, 1056 (2025).

12. Sugden, W. W. et al. Endoglin controls blood vessel diameter through endothelial cell shape changes in response to haemodynamic cues. Nat Cell Biol 19, 653–665 (2017).

13. Hill, R. A. et al. Regional Blood Flow in the Normal and Ischemic Brain Is Controlled by Arteriolar Smooth Muscle Cell Contractility and Not by Capillary Pericytes. Neuron 87, 95–110 (2015).

14. Sazonova, O. V. et al. Extracellular matrix presentation modulates vascular smooth muscle cell mechanotransduction. Matrix Biology 41, 36–43 (2015).

15. Rombouts, K. B. et al. The role of vascular smooth muscle cells in the development of aortic aneurysms and dissections. Eur J Clin Investigation 52, e13697 (2022).

16. Almaça, J., Weitz, J., Rodriguez-Diaz, R., Pereira, E. & Caicedo, A. The Pericyte of the Pancreatic Islet Regulates Capillary Diameter and Local Blood Flow. Cell Metabolism 27, 630–644.e4 (2018).

17. Bahrami, N. & Childs, S. J. Development of vascular regulation in the zebrafish embryo. Development 147, dev183061 (2020).

18. Hartmann, D. A. et al. Brain capillary pericytes exert a substantial but slow influence on blood flow. Nat Neurosci 24, 633–645 (2021).

19. Leonard, E. V. et al. Regenerating vascular mural cells in zebrafish fin blood vessels are not derived from pre-existing mural cells and differentially require Pdgfrb signalling for their development. Development 149, dev199640 (2022).

20. Ando, K. et al. Clarification of mural cell coverage of vascular endothelial cells by live imaging of zebrafish. Development dev.132654 (2016) doi:10.1242/dev.132654.

21. Attwell, D., Mishra, A., Hall, C. N., O’Farrell, F. M. & Dalkara, T. What is a pericyte? J Cereb Blood Flow Metab 36, 451–455 (2016).

22. Berthiaume, A.-A., Hartmann, D. A., Majesky, M. W., Bhat, N. R. & Shih, A. Y. Pericyte Structural Remodeling in Cerebrovascular Health and Homeostasis. Front. Aging Neurosci. 10, 210 (2018).

23. Figueiredo, A. M. et al. Phosphoinositide 3-Kinase–Regulated Pericyte Maturation Governs Vascular Remodeling. Circulation 142, 688–704 (2020).

24. Ando, K. et al. KCNJ8/ABCC9-containing K-ATP channel modulates brain vascular smooth muscle development and neurovascular coupling. Developmental Cell 57, 1383–1399.e7 (2022).

25. Vanlandewijck, M. et al. A molecular atlas of cell types and zonation in the brain vasculature. Nature 554, 475–480 (2018).

26. Murali, A. & Rajalingam, K. Small Rho GTPases in the control of cell shape and mobility. Cell. Mol. Life Sci. 71, 1703–1721 (2014).

27. Álvarez-Aznar, A. et al. Cdc42 is crucial for mural cell migration, proliferation and patterning of the retinal vasculature. Vascular Pharmacology 159, 107472 (2025).

28. Montanez, E., Wickström, S. A., Altstätter, J., Chu, H. & Fässler, R. α-parvin controls vascular mural cell recruitment to vessel wall by regulating RhoA/ROCK signalling. EMBO J 28, 3132–3144 (2009).

29. Gore, A. V., Monzo, K., Cha, Y. R., Pan, W. & Weinstein, B. M. Vascular Development in the Zebrafish. Cold Spring Harbor Perspectives in Medicine 2, a006684–a006684 (2012).

30. Zeng, H. Mural-VISTA: a tool for mural cell-vessel interaction assessment and multiscale single-cell topo-morphological analysis.

31. Sur, A. et al. Single-cell analysis of shared signatures and transcriptional diversity during zebrafish development. Developmental Cell 58, 3028–3047.e12 (2023).

32. Stratman, A. N. et al. Mural-Endothelial cell-cell interactions stabilize the developing zebrafish dorsal aorta. Development dev.143131 (2016) doi:10.1242/dev.143131.

33. Shih, Y.-H., Portman, D., Idrizi, F., Grosse, A. & Lawson, N. D. Integrated molecular analysis identifies a conserved pericyte gene signature in zebrafish. Development 148, dev200189 (2021).

34. Lancino, M. et al. Anisotropic organization of circumferential actomyosin characterizes hematopoietic stem cells emergence in the zebrafish. eLife 7, e37355 (2018).

35. Somlyo, A. P. & Somlyo, A. V. Ca^2+^ Sensitivity of Smooth Muscle and Nonmuscle Myosin II: Modulated by G Proteins, Kinases, and Myosin Phosphatase. Physiological Reviews 83, 1325–1358 (2003).

36. Touyz, R. M. et al. Vascular smooth muscle contraction in hypertension. Cardiovascular Research 114, 529–539 (2018).

37. Amano, M., Nakayama, M. & Kaibuchi, K. Rho-kinase/ROCK: A key regulator of the cytoskeleton and cell polarity. Cytoskeleton 67, 545–554 (2010).

38. Hall, C. N. et al. Capillary pericytes regulate cerebral blood flow in health and disease. Nature 508, 55–60 (2014).

39. Cheng, S. et al. Hemodynamics regulate spatiotemporal artery muscularization in the developing circle of Willis. eLife 13, RP94094 (2024).

40. Isogai, S., Horiguchi, M. & Weinstein, B. M. The Vascular Anatomy of the Developing Zebrafish: An Atlas of Embryonic and Early Larval Development. Developmental Biology 230, 278–301 (2001).

41. Hartmann, D. A. et al. Pericyte structure and distribution in the cerebral cortex revealed by high-resolution imaging of transgenic mice. Neurophoton 2, 041402 (2015).

42. Berthiaume, A.-A. et al. Dynamic Remodeling of Pericytes In Vivo Maintains Capillary Coverage in the Adult Mouse Brain. Cell Reports 22, 8–16 (2018).

43. Dessalles, C. A., Babataheri, A. & Barakat, A. I. Pericyte mechanics and mechanobiology. Journal of Cell Science 134, jcs240226 (2021).

44. Rajan, A. M., Ma, R. C., Kocha, K. M., Zhang, D. J. & Huang, P. Dual function of perivascular fibroblasts in vascular stabilization in zebrafish. PLoS Genet 16, e1008800 (2020).

45. Loirand, G., Guérin, P. & Pacaud, P. Rho Kinases in Cardiovascular Physiology and Pathophysiology. Circulation Research 98, 322–334 (2006).

46. Morooka, N. et al. Angpt1 binding to Tie1 regulates the signaling required for lymphatic vessel development in zebrafish. Development 151, dev202269 (2024).

47. Jiang, Z., Carlantoni, C., Allanki, S., Ebersberger, I. & Stainier, D. Y. R. Tek (Tie2) is not required for cardiovascular development in zebrafish. Development 147, dev193029 (2020).

48. Gjini, E. et al. Zebrafish Tie-2 shares a redundant role with Tie-1 in heart development and regulates vessel integrity. Disease Models & Mechanisms 4, 57–66 (2011).

49. Crestani, S., Webb, R. C. & Da Silva-Santos, J. E. High-Salt Intake Augments the Activity of the RhoA/ROCK Pathway and Reduces Intracellular Calcium in Arteries From Rats. American Journal of Hypertension 30, 389–399 (2017).

50. Loirand, G. & Pacaud, P. Involvement of Rho GTPases and their regulators in the pathogenesis of hypertension. Small GTPases 5, e983866 (2014).

51. Kimmel, C. B., Ballard, W. W., Kimmel, S. R., Ullmann, B. & Schilling, T. F. Stages of embryonic development of the zebrafish. Developmental Dynamics 203, 253–310 (1995).

52. Jin, S.-W., Beis, D., Mitchell, T., Chen, J.-N. & Stainier, D. Y. R. Cellular and molecular analyses of vascular tube and lumen formation in zebrafish. Development 132, 5199–5209 (2005).

53. Phng, L.-K., Stanchi, F. & Gerhardt, H. Filopodia are dispensable for endothelial tip cell guidance. Development 140, 4031–4040 (2013).

54. Fukuhara, S. et al. Visualizing the cell-cycle progression of endothelial cells in zebrafish. Developmental Biology 393, 10–23 (2014).

55. Antiga, L. & Steinman, D. A. Robust and Objective Decomposition and Mapping of Bifurcating Vessels. IEEE Trans. Med. Imaging 23, 704–713 (2004).

56. Fujita, Y., Nakamura, Y. & Shiller, Z. Dual Dijkstra Search for paths with different topologies. in 2003 IEEE International Conference on Robotics and Automation (Cat. No.03CH37422) vol. 3 3359–3364 (IEEE, Taipei, Taiwan, 2003).

57. Baltaci, O. F. et al. Vascular mural cells protect the adult brain from haemorrhage but do not control the blood-brain barrier in developing zebrafish. Preprint at 10.7554/eLife.104061.1 (2025).

