## Supplemental Figure for "Mural cell contractility regulates vessel diameter by controlling cell morphology and vessel coverage"

Supplementary Figure 1

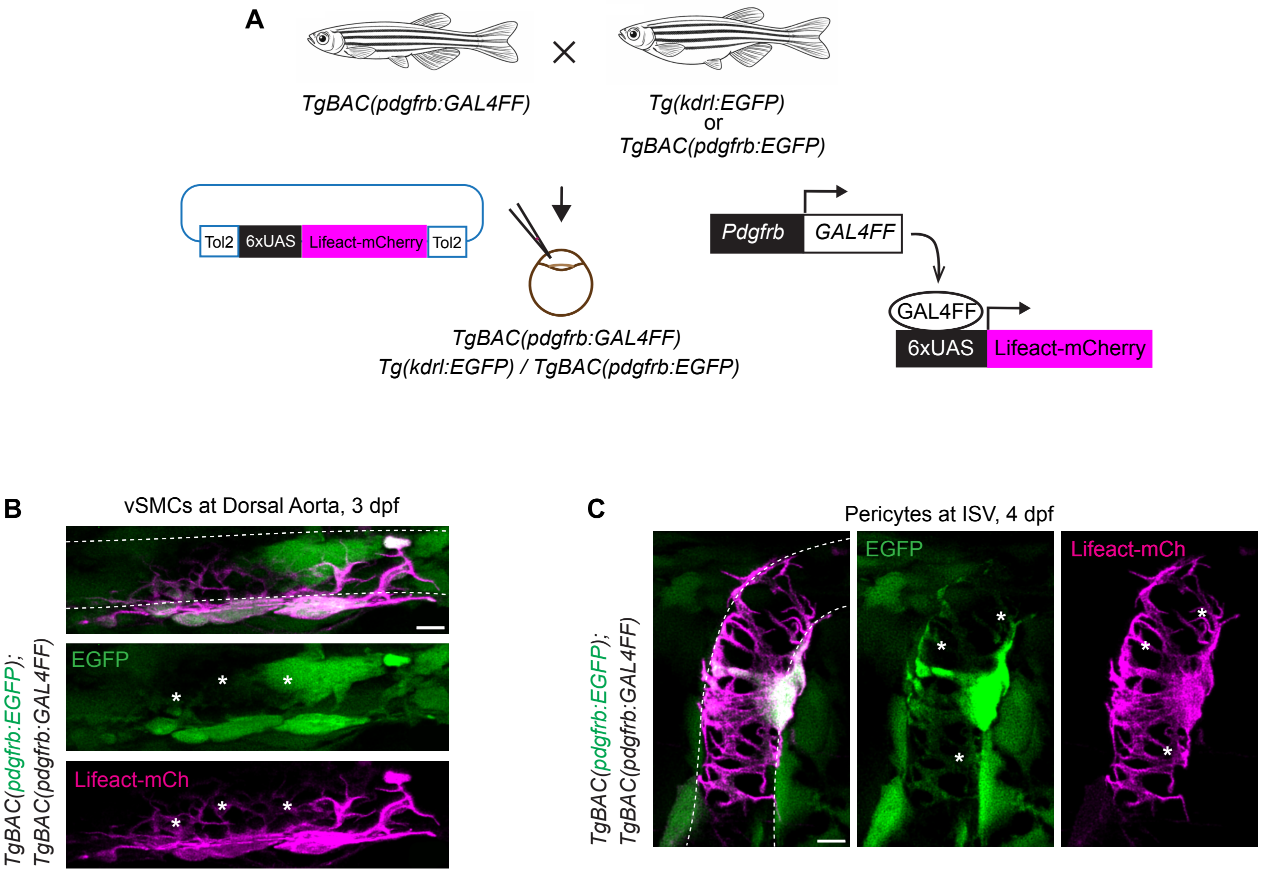


**Supplementary Figure 1** GAL4FF/UAS-driven Lifeact–mCherry enables high-resolution visualisation of mural cell morphology in zebrafish.

**A** Experimental strategy to label mural cells using the GAL4FF/UAS system. *TgBAC(pdgfrb:GAL4FF)* zebrafish were crossed with *Tg(kdrl:EGFP)* line to label endothelial cells or *TgBAC(pdgfrb:EGFP)* line to label mural cells. Embryos were injected with a 6×UAS:Lifeact–mCherry construct, resulting in *pdgfrb*-driven GAL4FF activation of Lifeact–mCherry expression in mural cells. **B – C** High-magnification confocal z-stacks of vSMCs and pericytes along the DA (**B**) and ISV (**C**) at 3 dpf. GFP labels mural cells, while Lifeact–mCherry highlights actin-rich cell bodies and processes, revealing fine processes not readily resolved by GFP alone. Scale bars, 5 μm. DA dorsal aorta, ISV intersegmental vessel, aISV arterial ISV, vISV venous ISV.

Supplementary Figure 2

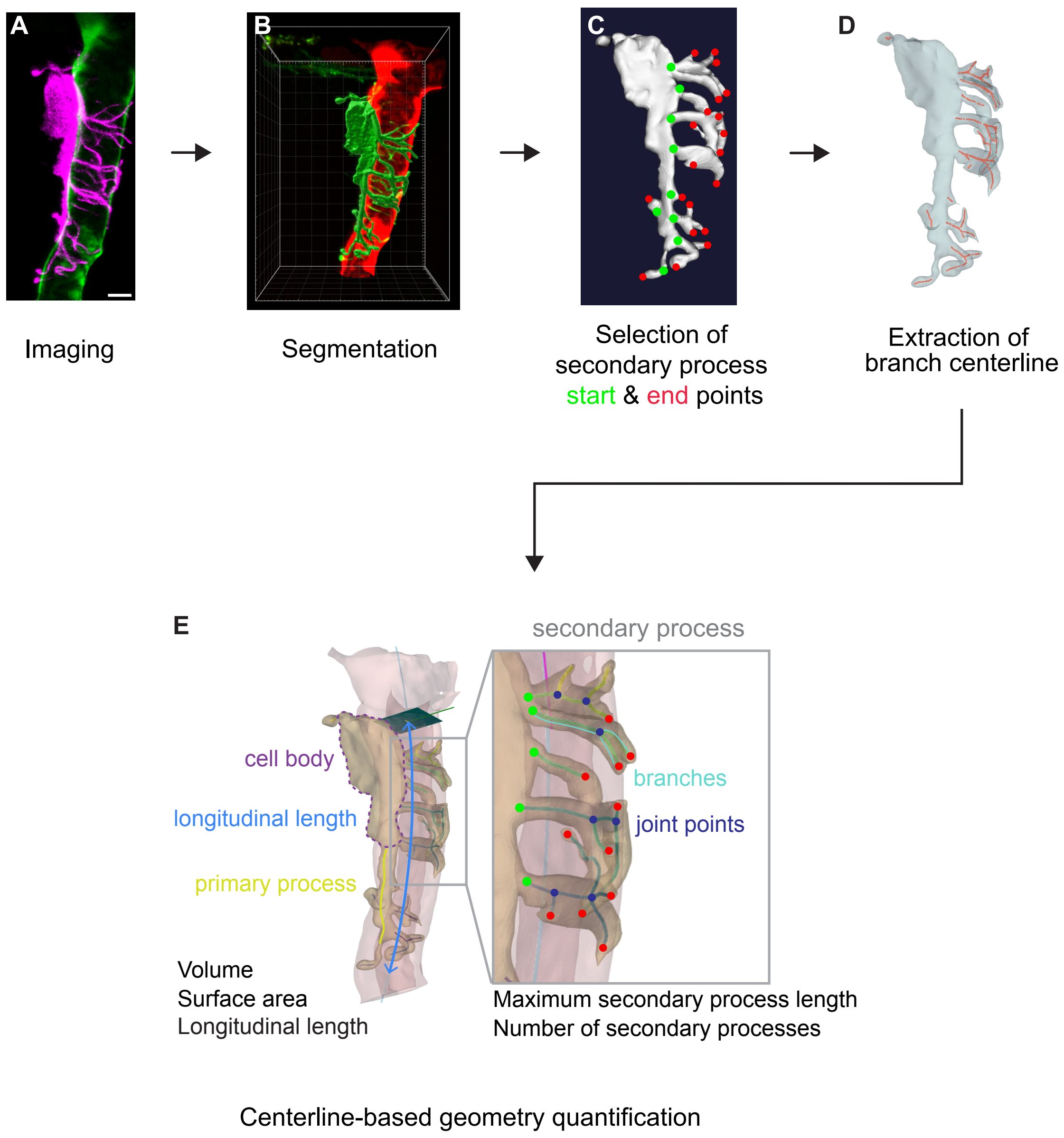


**Supplementary Figure 2** Workflow for 3D reconstruction and morphometric quantification of mural cell geometry. **A** Maximum intensity projection of confocal z-stacks of a pericyte labelled with Lifeact-mCherry. **B** Three-dimensional segmentation of the mural cell from confocal z-stacks, separating the cell volume from background signal. **C** Identification of secondary processes with annotation of start points (green) and end points (red). **D** Extraction of branch centrelines from the segmented cell volume to generate a skeletonised representation of cellular processes. **E** Quantitative analysis based on centreline geometry. Inset indicates higher magnification views of secondary process structure.

Supplementary Figure 3

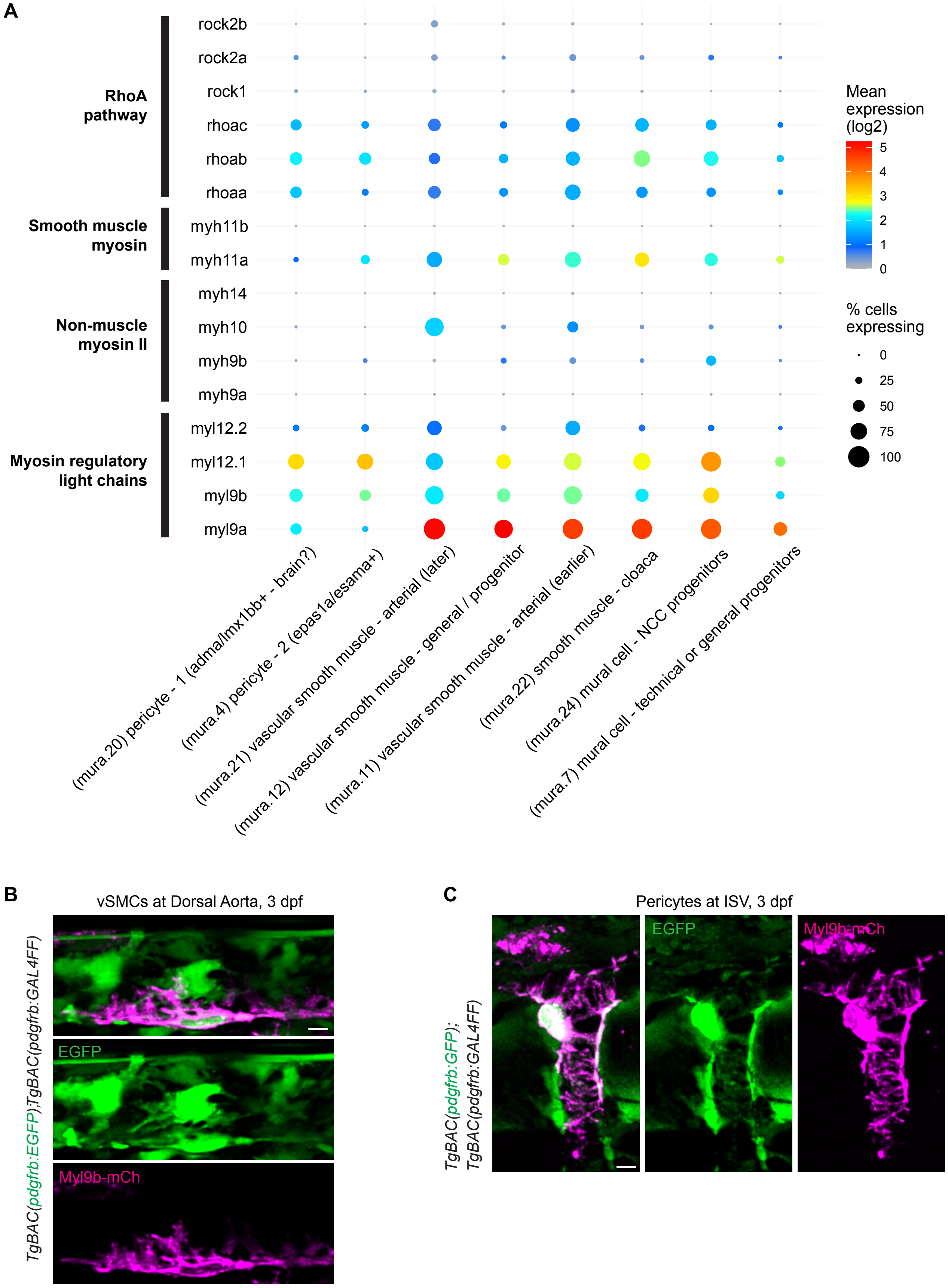


**Supplementary Figure 3** Expression and localisation of actomyosin regulatory components in zebrafish mural cells. **A** Dot plot showing the expression of genes associated with the RhoA pathway, smooth muscle myosin, non-muscle myosin II and myosin regulatory light chains across zebrafish pericyte and vSMC clusters. Dot size indicates the percentage of cells expressing each gene, and colour indicates mean expression level (log2). Data were obtained from Daniocell (https://daniocell.nichd.nih.gov). **B – C** Maximum intensity projection of confocal z-stacks of vSMCs (**B**) and pericytes (**C**) along the DA and ISV at 3 dpf in *TgBAC(pdgfrb:EGFP); TgBAC(pdgfrb:GAL4FF)* embryos expressing Myl9b–mCherry. Myl9b–mCherry is enriched in the soma and extends into primary and secondary processes. Scale bars, 5 µm.

Supplementary Figure 4

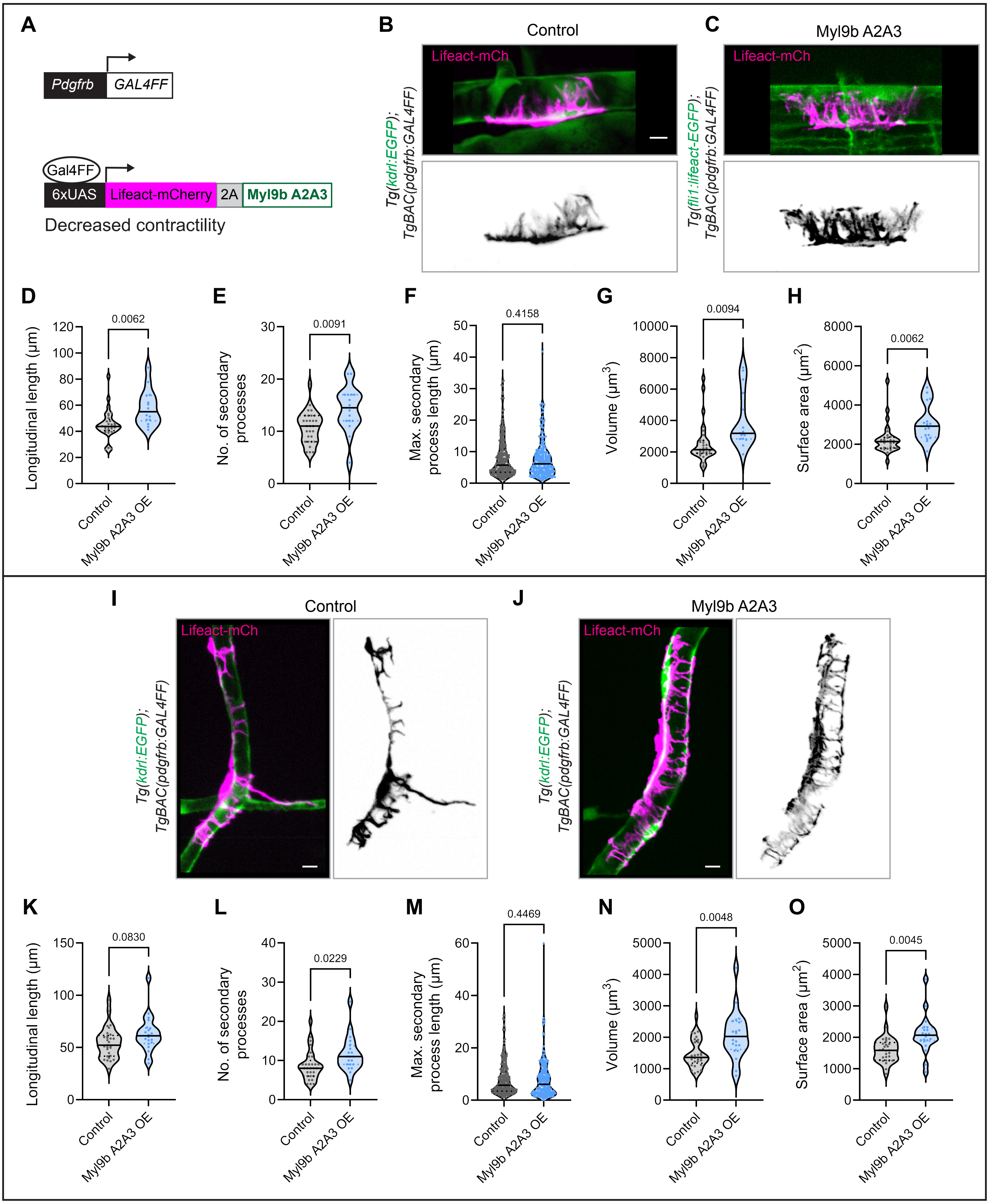


**Supplementary Figure 4** Decreased Myl9b activity promotes mural cell enlargement and process extension.
**A** Schematic of the GAL4FF/UAS strategy used to express Myl9bA2A3 in mural cells. Myl9b A2A3 was expressed under the control of the *pdgfrb:Gal4FF* driver together with Lifeact-mCherry, resulting in reduced contractility. **B – C** Representative images of control (**B**) and Myl9bA2A3 - overexpressing (**C**) vSMCs at 5 dpf, showing Lifeact-mCherry labelled vSMC morphology. **D – H** Quantification of vSMC longitudinal length (**D**), number of secondary processes (**E**), maximum secondary process length (**F**), cell volume (**G**) and surface area (**H**). **I – J** Representative images of control (**I**) and Myl9bA2A3-overexpressing (**J**) pericytes. **K – O** Quantification of pericyte longitudinal length (**K**), number of secondary processes (**L**), maximum secondary process length (**M**), cell volume (**N**) and surface area (**O**). Myl9bA2A3 overexpression increases cell size and promotes extension of cellular processes in both vSMCs and pericytes. Statistical comparisons and P values are indicated in the graphs. All data are collected from 2 independent experiments and analyzed by Welch's t test.

Supplementary Figure 5

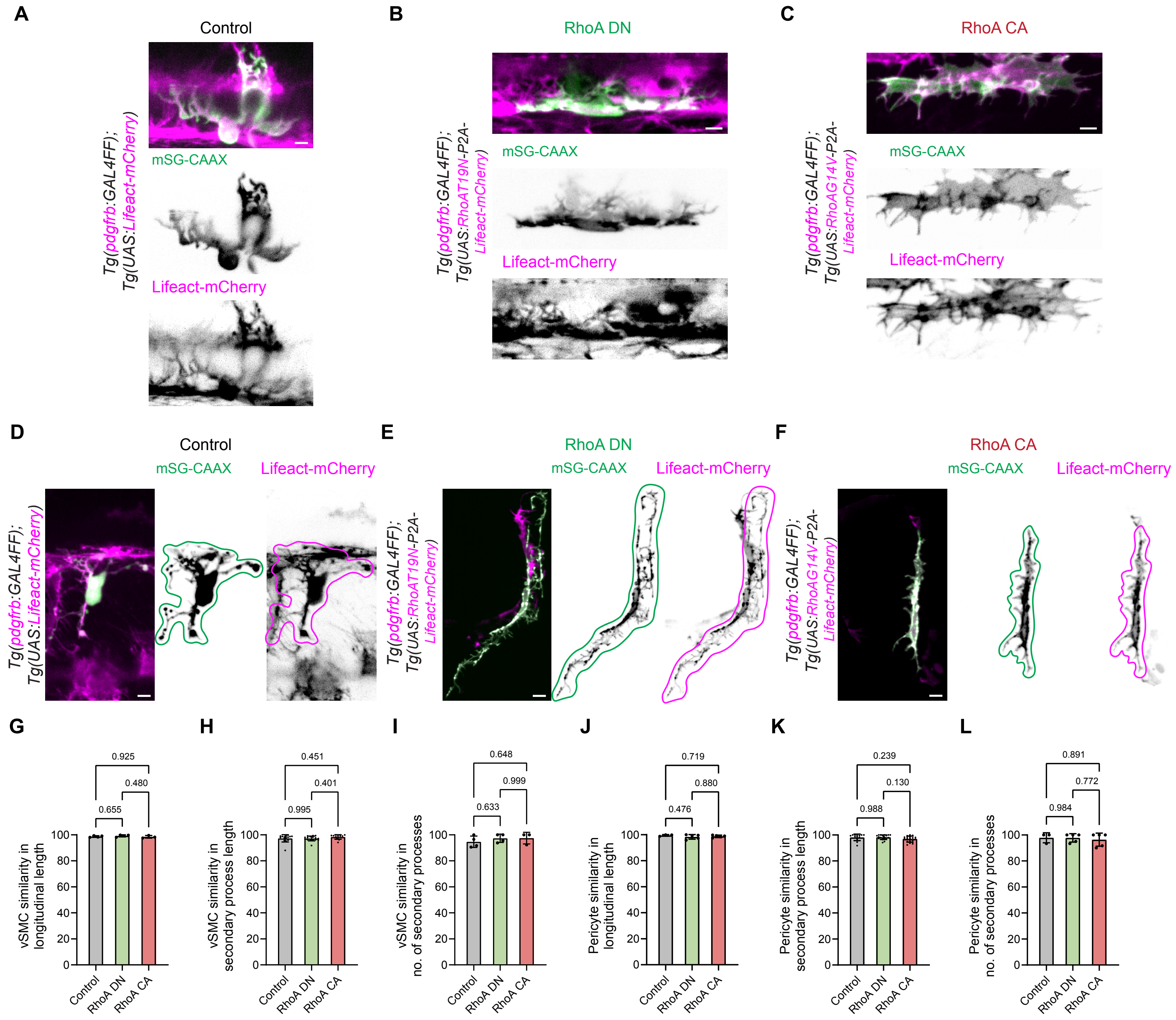


**Supplementary Figure 5** Validation of Lifeact-mCherry-based mural cell morphometrics using mStayGold-CAAX. **A – C** Representative images of vSMCs expressing membrane-localised mStayGold-CAAX (green) and Lifeact-mCherry (magenta) in control (**A**), *Tg(RhoA DN-OE)* (**B**) and *Tg(RhoA CA-OE)* (**C**) embryos at 5 dpf. Images show comparable labelling patterns and cellular boundaries between mStayGold-CAAX and Lifeact-mCherry across all three conditions. **D – F** Representative images of individual pericytes used for morphometric analysis, with cell outlines independently traced based on mStayGold-CAAX (green) and Lifeact-mCherry (magenta) signals in control (**D**), *Tg(RhoA DN-OE)* (**E**) and *Tg(RhoA CA-OE)* (**F**) embryos at 5 dpf. **G – I** Quantification of the similarity between mStayGold-CAAX- and Lifeact-mCherry-based measurements of vSMC longitudinal length (**G**), secondary process length (**H**) and number of secondary processes (**I**). **J – L** Quantification of the similarity between mStayGold-CAAX- and Lifeact-mCherry-based measurements of pericyte longitudinal length (**J**), secondary process length (**K**) and number of secondary processes (**L**). Similarity was calculated for each measurement as Lifeact-mCherry-based measurement divided by the corresponding mStayGold-CAAX-based measurement × 100%. Values close to 100% indicate close overlap between the two labelling approaches. Across all measured parameters, Lifeact-mCherry-based measurements closely matched those obtained using mStayGold-CAAX in control, *Tg(RhoA DN-OE)* and *Tg(RhoA CA-OE)* embryos. Statistical significance was assessed by one-way ANOVA with multiple-comparisons testing. Scale bars, 5 μm.

Supplementary Figure 6

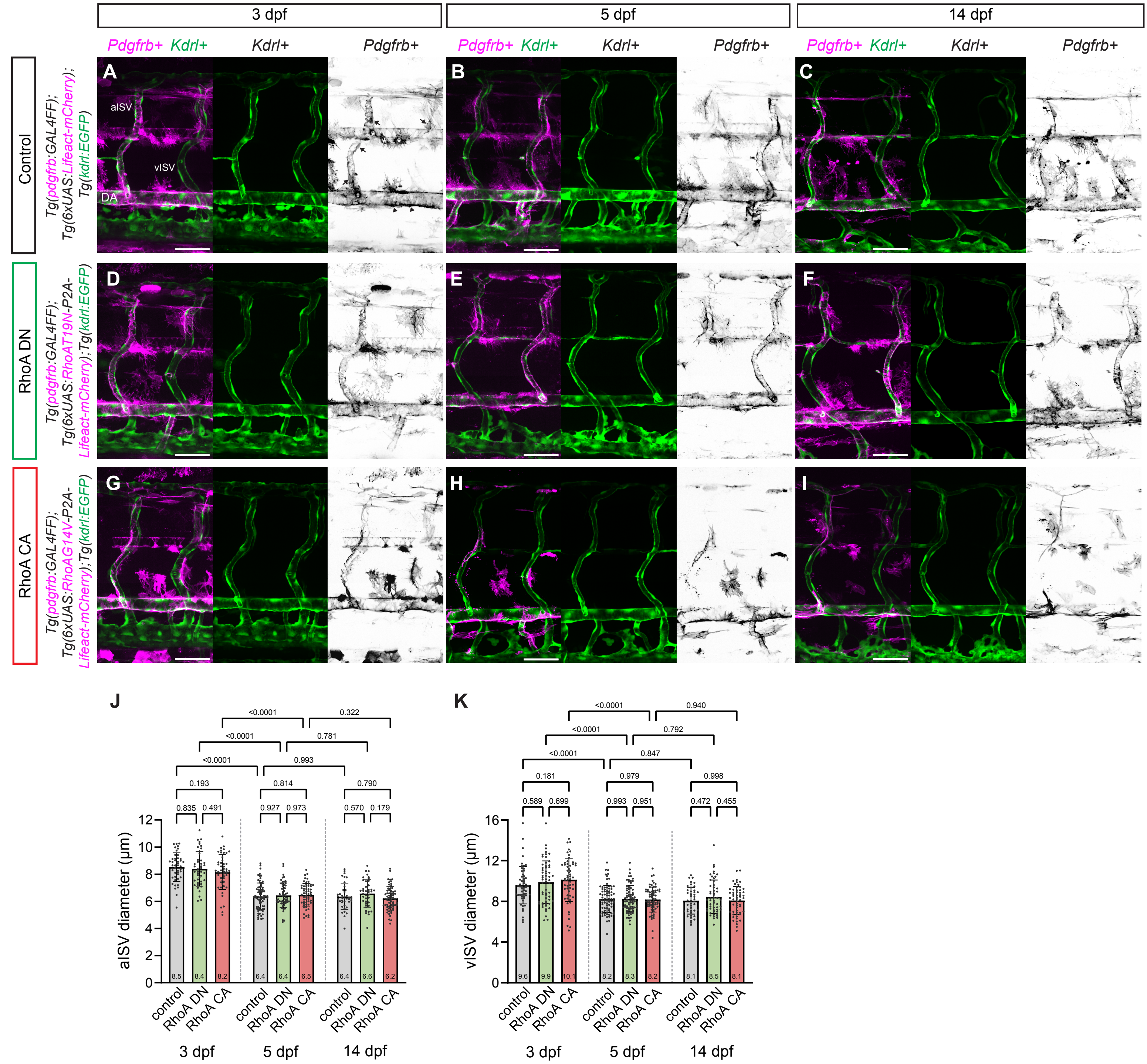


**Supplementary Figure 6** Pericyte contractility does not affect intersegmental vessel formation and remodelling.
**A – C** Maximum intensity projection of confocal z-stacks of trunk vasculature in *Tg(kdrl:EGFP); TgBAC(pdgfrb:GAL4FF); Tg(6×UAS:Lifeact–mCherry)* embryos at 3, 5 and 14 dpf. Magenta, *Pdgfrb*+ mural cells; green, *Kdrl*+ endothelial cells. **D – F** Representative confocal z-stacks of trunk vasculature in *Tg(kdrl:EGFP); TgBAC(pdgfrb:GAL4FF); Tg(6×UAS:RhoA T19N–P2A–Lifeact–mCherry)* embryos at corresponding developmental stages. **G – I** Representative confocal z-stacks of trunk vasculature in *Tg(kdrl:EGFP); TgBAC(pdgfrb:GAL4FF); Tg(6×UAS:RhoA G14V–P2A–Lifeact–mCherry)* embryos at corresponding developmental stages. **J – K** Quantification of aISV and vISV diameters at 3, 5 and 14 dpf under control, RhoA DN and RhoA CA conditions. Control: n= 44/75/40 aISVs and 55/80/43 vISVs from 13/21/13 embryos at 3/5/14 dpf; RhoA DN: n= 43/56/40 aISVs and 58/72/53 vISVs from 13/17/13 embryos at 3/5/14 dpf; RhoA CA: n= 49/58/26 aISVs and 56/69/26 vISVs from 14/16/21 embryos at 3/5/14 dpf. ISV diameters are collected from 2 independent experiments and analyzed by two-way ANOVA with Tukey's multiple comparisons test. Scale bars, 50 µm. DA dorsal aorta, ISV intersegmental vessel, aISV arterial ISV, vISV venous ISV.

Supplementary Figure 7

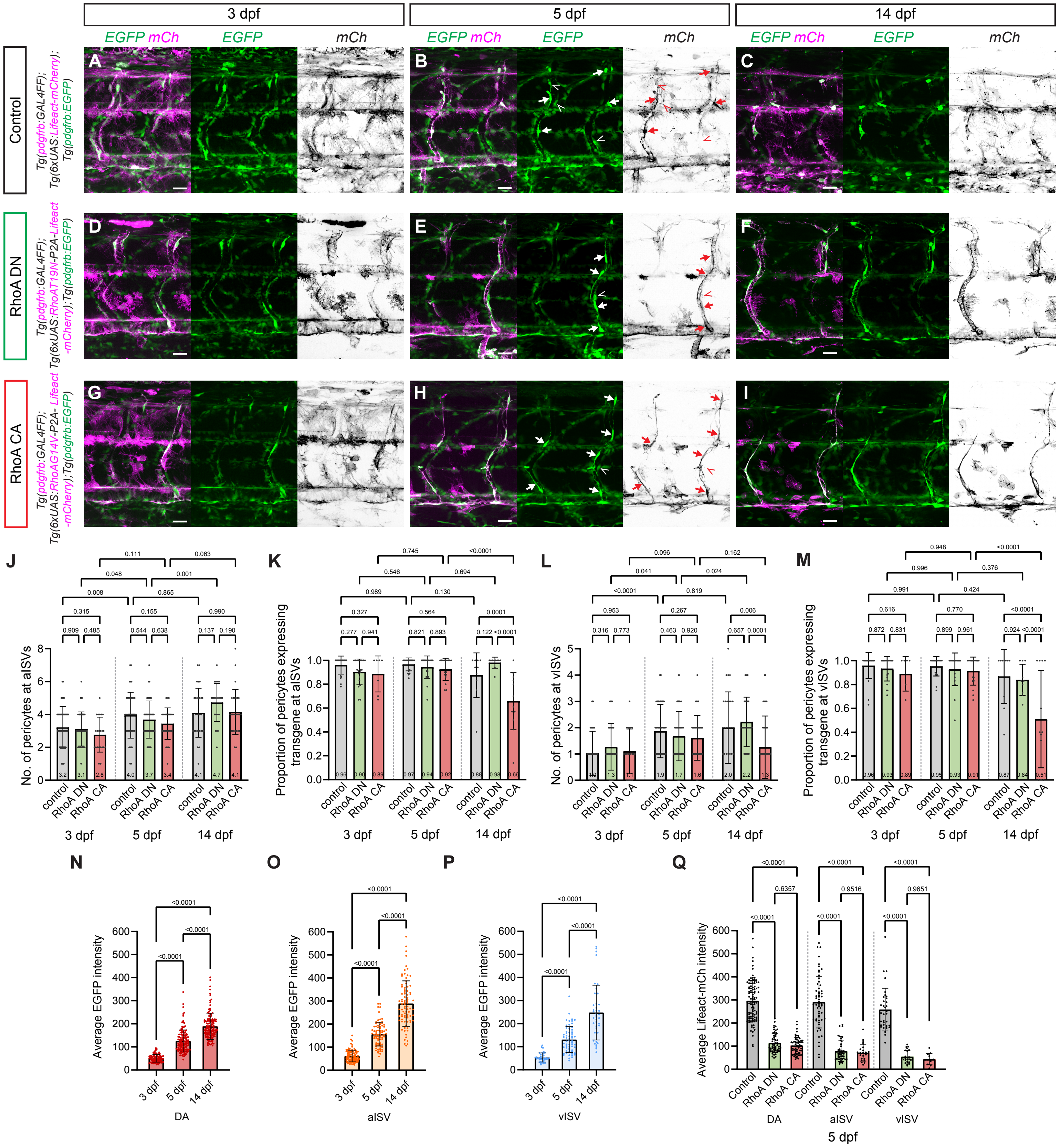


**Supplementary Figure 7** Developmental changes in mural cell and perivascular fibroblast abundance and transgene expression. **A – C** Representative maximum intensity projections of confocal z-stacks showing the trunk vasculature in *TgBAC(pdgfrb:EGFP); TgBAC(pdgfrb:GAL4FF); Tg(6×UAS:Lifeact–mCherry)* embryos at 3, 5 and 14 dpf. Green, mural cells labelled by *pdgrfb*:EGFP; magenta/black, mural cells expressing transgene using GAL4FF/UAS strategy. Arrows indicate pericytes; arrowheads indicate perivascular fibroblasts. **D – F** Representative maximum intensity projections of confocal z-stacks showing the trunk vasculature in *TgBAC(pdgfrb:EGFP); TgBAC(pdgfrb:GAL4FF); Tg(6×UAS:RhoA T19N–P2A–Lifeact–mCherry)* embryos at corresponding developmental stages. **G – I** Representative maximum intensity projections of confocal z-stacks showing the trunk vasculature in *TgBAC(pdgfrb:EGFP); TgBAC(pdgfrb:GAL4FF); Tg(6×UAS:RhoA G14V–P2A–Lifeact–mCherry)* embryos at corresponding developmental stages. **J – M** Number of pericytes associated with ISVs (**J**, **L**) and the proportion of pericytes expressing the transgene relative to the total number of pericytes (**K**, **M**) at 3, 5 and 14 dpf. **N – P** Average *pdgfrb*:EGFP fluorescence intensity in the DA (**N**), aISVs (**O**) and vISVs (**P**) during development of control embryos. **Q** Average Lifeact-mCherry fluorescence intensity in the DA, aISVs and vISVs of 5 dpf control, *Tg(RhoA DN-OE)* and *Tg(RhoA CA-OE)* embryos. Data are presented as mean ± SD; individual data points are shown. Statistical comparisons and P values are indicated in the graphs. Scale bars, 5 μm. All data are collected from 2 independent experiments and analyzed by two-way ANOVA with Tukey's multiple comparisons test and ordinary one-way ANOVA with Tukey's multiple comparisons test.

Supplementary Figure 8

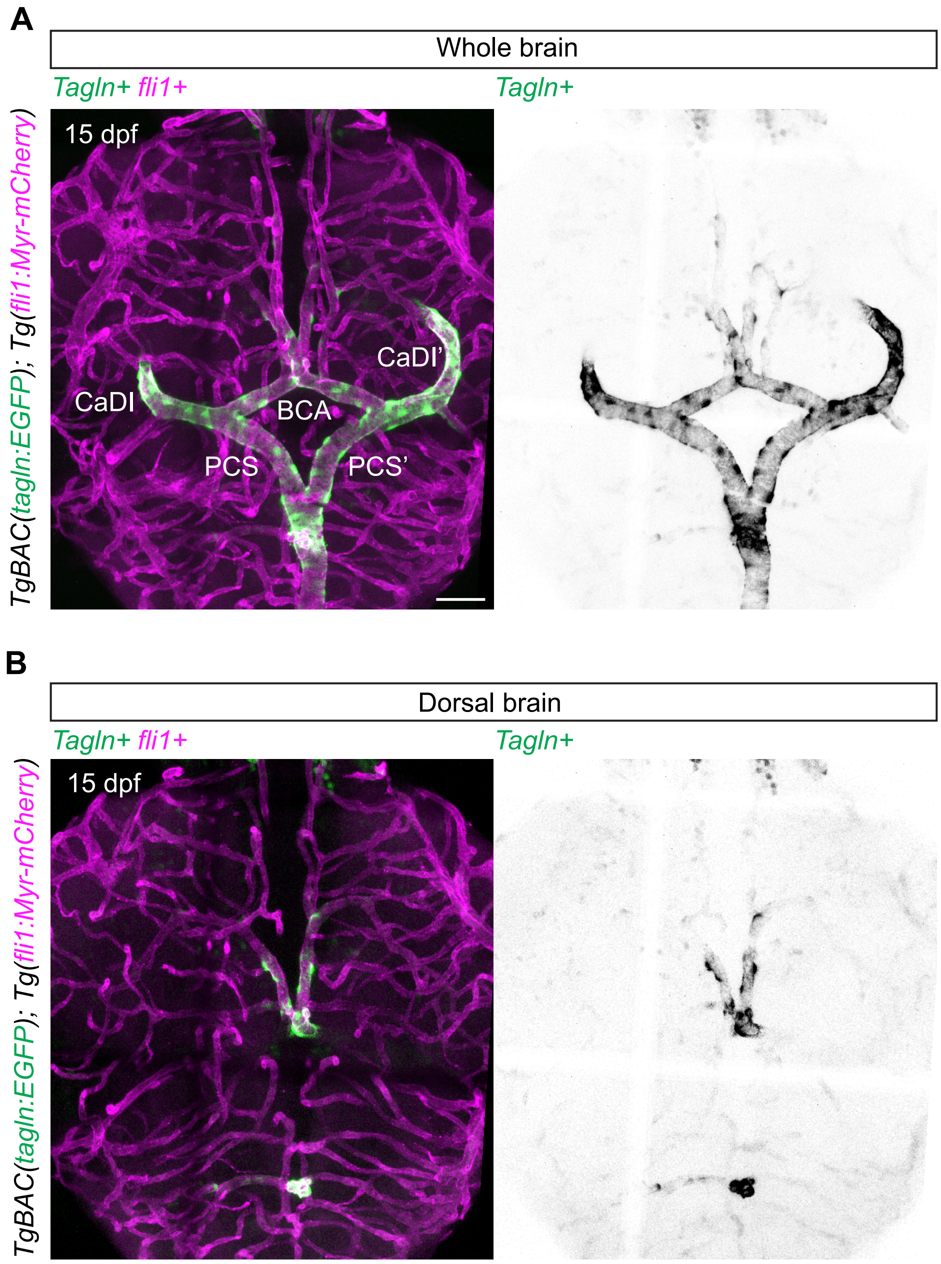


**Supplementary Figure 8** Distinct Tagln expression in vSMCs of the brain vasculature.
**A** Representative confocal z-stacks of the whole brain at 15 dpf in *TgBAC(tagln:EGFP); Tg(fli1:myr–mCherry)* embryos, showing differentiated vSMCs (*Tagln*+, green) and endothelial cells (*fli1*+, magenta). Tagln expression is enriched in major vessels of the CoW. Right, Tagln signal shown alone. **B** Representative confocal z-stacks of the dorsal brain region at 15 dpf showing restricted Tagln expression in major vessels, with minimal signal detected in smaller calibre vessels. Right, Tagln signal shown alone. Scale Bar, 50 µm. CoW Circle of Willis, BCA basal communicating artery, CaDI caudal division of the internal carotid artery, PCS posterior communicating segment.

Supplementary Figure 9

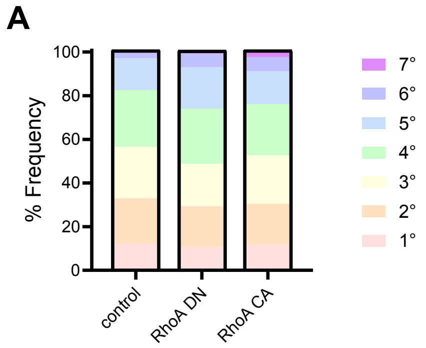


**Supplementary Figure 9** Mural cell contractility regulates brain vascular branching.
**A** Quantification of CtA branch diameters across different branch orders in 14 dpf control, RhoA DN and RhoA CA embryos. Control: n = 246 section vessels from 6 embryos, RhoA DN; n = 297 section vessels from 8 embryos, RhoA CA: n = 367 section vessels from 9 embryos. Control vs RhoA DN, 7° vs other orders, Fisher's exact test, p = 0.5035; control vs RhoA CA, 7° vs other orders, Fisher's exact test, p= 0.0132.

Supplementary Figure 10

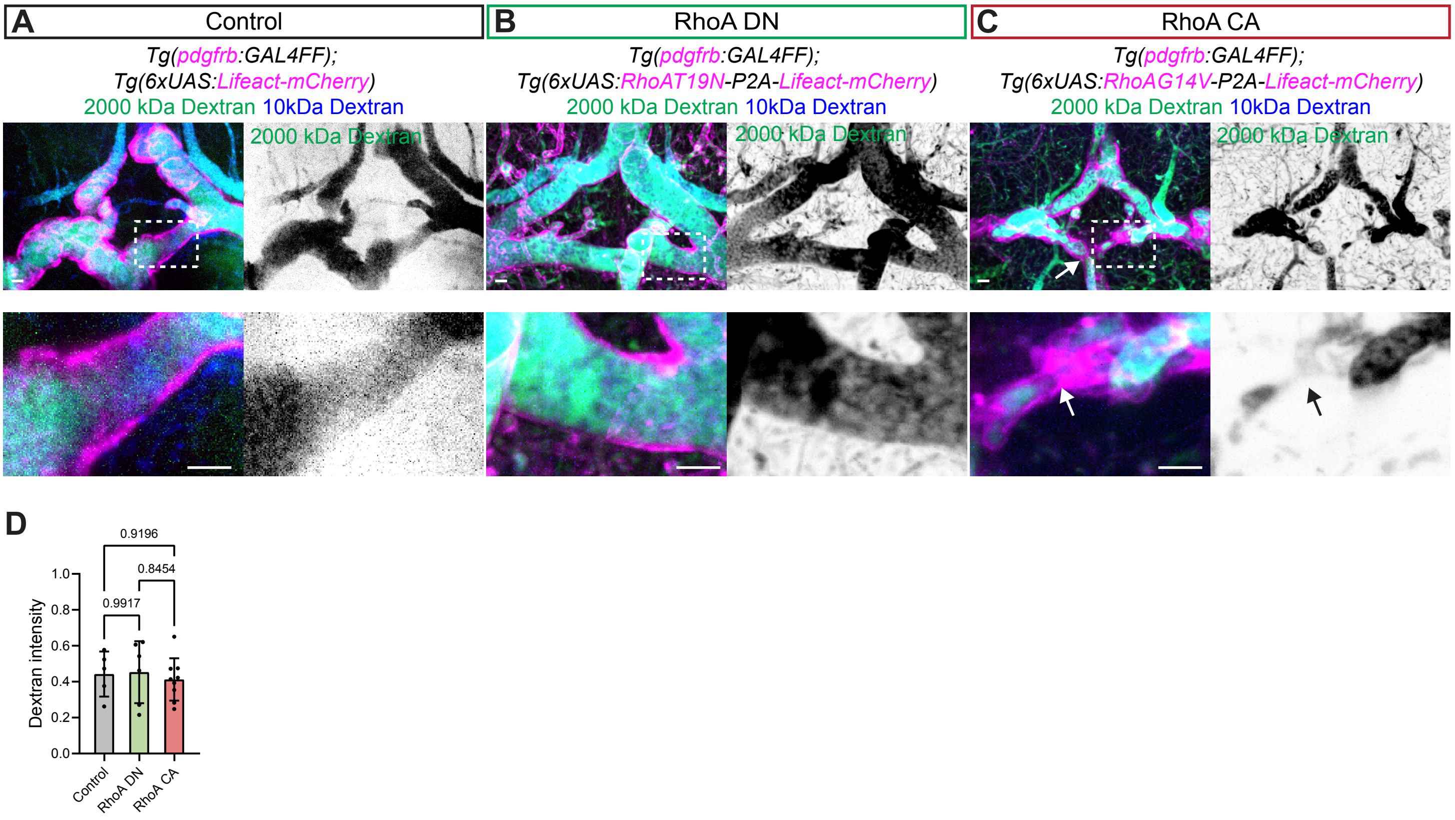


**Supplementary Figure 10** Increased mural cell contractility causes stenosis of CoW vessels in adult zebrafish .
**A – C** Representative images of CoW vessels from adult zebrafish expressing Lifeact-mCherry in mural cells of control (**A**), RhoA DN (**B**) and RhoA CA (**C**) zebrafish. 2000 kDa dextran was used to visualise the vascular lumen, while 10 kDa dextran was used as a smaller molecular-weight tracer for tracer leakage quantification. Enlarged views of the boxed regions are shown . Arrows indicate regions of vessel narrowing/stenosis. **D** Fluorescent tracer assays in 10-month-old (control) and 13-month-old (RhoA DN and RhoA CA) brain. 10 kDa Dextran–Cascade Blue was used to detect tracer extravasation to the brain parenchyma and were coinjected with 2000 kDa Dextran Fluorescein to normalize vascular tracer intensity (n= 5 control; 6 RhoA DN; 8 RhoA CA zebrafish). All data were collected from 2 independent experiments and analysed by one-way ANOVA with Tukey's multiple comparisons test.

**Video 1** Control pericyte shows progressive retraction of lamellar secondary process.
Time-lapse imaging of lamellar process dynamics in trunk pericytes of *TgBAC(pdgfrb:GAL4FF); Tg(6×UAS:Lifeact–mCherry)* embryos from 80 to 90 hpf. Lifeact-mCherry labels F-actin in pericytes, revealing dynamic extension and retraction of lamellar processes along the vessel wall. Time interval, 20 min; total duration, 580 min. Scale bar, 5 μm.

**Video 2** RhoA DN–overexpressing pericytes show minimal lamellar process retraction.
Time-lapse imaging of lamellar process dynamics in trunk pericytes of *TgBAC(pdgfrb:GAL4FF); Tg(6×UAS:RhoA T19N–P2A–Lifeact–mCherry)* embryos from 80 to 90 hpf. Lifeact-mCherry labels F-actin in pericytes, revealing that RhoA DN–overexpressing pericytes show minimal change in the length of lamellar secondary processes between 80 and 90 hpf. Time interval: 20 minutes. Time interval, 20 min; total duration, 580 min. Scale bar, 5 μm.

**Video 3** RhoA CA–overexpressing pericyte accelerates shortening of lamellar secondary processes.
Time-lapse imaging of lamellar process dynamics in trunk pericytes of *TgBAC(pdgfrb:GAL4FF); Tg(6×UAS:RhoA G14V–P2A–Lifeact–mCherry)* embryos from 80 to 90 hpf. Increased RhoA activity induces earlier shortening of lamellar processes. Red line delineates the shape of the pericyte on the first frame of the movie. Time interval, 20 min; total duration, 580 min. Scale bar, 5 μm.

**Video 4** Cyclic deformation of the DA in control zebrafish at 84–85 hpf.
Time-lapse imaging of the DA shows rhythmic diameter changes driven by pulsatile blood flow in *Tg(kdrl:EGFP); TgBAC(pdgfrb:GAL4FF); Tg(6×UAS:Lifeact–mCherry)* embryos at 84–85 hpf. The ventral DA wall undergoes cyclic expansion and relaxation during cardiac cycle, reflecting high elasticity of the vessel wall at this developmental stage. Time interval, 10 ms; total duration, 3 s. Scale bar, 5 μm.

**Video 5** Increased DA wall displacement in *Tg(RhoA DN-OE)* zebrafish.
Time-lapse imaging reveals enhanced cyclic deformation of the ventral DA wall in *Tg(kdrl:EGFP); TgBAC(pdgfrb:GAL4FF); Tg(6×UAS:RhoA T19N–P2A–Lifeact–mCherry)* embryos at 84–85 hpf. The increased peak-to-trough displacement indicates elevated vessel wall compliance in response to pulsatile pressure, consistent with reduced mural cell contractility. Time interval, 10 ms; total duration, 3 s. Scale bar, 5 μm.

**Video 6** Reduced DA wall displacement in *Tg(RhoA CA-OE)* zebrafish.
Time-lapse imaging shows undetectable cyclic deformation of the ventral DA wall in *Tg(kdrl:EGFP); TgBAC(pdgfrb:GAL4FF); Tg(6×UAS:RhoA G14V–P2A–Lifeact–mCherry)* embryos at 84–85 hpf. The decreased peak-to-trough displacement reflects reduced vessel wall compliance and increased stiffness, consistent with elevated mural cell contractility. Time interval, 10 ms; total duration, 3 s. Scale bar, 5 μm.
